# Capturing Brain-Cognition Relationship: Integrating Task-Based fMRI Across Tasks Markedly Boosts Prediction and Test-Retest Reliability

**DOI:** 10.1101/2021.10.31.466638

**Authors:** Alina Tetereva, Jean Li, Jeremiah D. Deng, Argyris Stringaris, Narun Pat

## Abstract

Capturing individual differences in cognition is central to human neuroscience. Yet our ability to estimate cognitive abilities via brain MRI is still poor in both prediction and reliability. Our study tested if this inability can be improved by integrating MRI signals across the whole brain and across modalities, including task-based functional MRI (tfMRI) of different tasks along with other non-task MRI modalities, such as structural MRI, resting-state functional connectivity. Using the Human Connectome Project (n=873, 473 females, after quality control), we directly compared predictive models comprising different sets of MRI modalities (e.g., seven tasks vs. non-task modalities). We applied two approaches to integrate multimodal MRI, stacked vs. flat models, and implemented 16 combinations of machine-learning algorithms. The stacked model integrating all modalities via stacking Elastic Net provided the best prediction (*r*=.57), relatively to other models tested, as well as excellent test-retest reliability (ICC=^~^.85) in capturing general cognitive abilities. Importantly, compared to the stacked model integrating across non-task modalities (*r*=.27), the stacked model integrating tfMRI across tasks led to significantly higher prediction (*r*=.56) while still providing excellent test-retest reliability (ICC=^~^.83). The stacked model integrating tfMRI across tasks was driven by frontal and parietal areas and by tasks that are cognition-related (working-memory, relational processing, and language). This result is consistent with the parieto-frontal integration theory of intelligence. Accordingly, our results contradict the recently popular notion that tfMRI is not reliable enough to capture individual differences in cognition. Instead, our study suggests that tfMRI, when used appropriately (i.e., by drawing information across the whole brain and across tasks and by integrating with other modalities), provides predictive and reliable sources of information for individual differences in cognitive abilities, more so than non-task modalities.

**Highlights:**

- Non-task MRI (sMRI, rs-fMRI) are often used for the brain-cognition relationship.
- Task-based fMRI has been deemed unreliable for capturing individual differences.
- We tested if drawing task-based fMRI information across regions/tasks improves prediction and reliability of the brain-cognition relationship.
- Our approach boosts prediction of task-based fMRI over non-task MRI.
- Our approach renders task-based fMRI reliable over time.
- Our approach shows the importance of the fronto-parietal areas in cognition.

## 1. Introduction

Relating individual differences in cognitive abilities to the brain has been focal to human neuroscience (Deary et al., 2010). Yet, we still cannot use brain data to capture individual differences in cognitive abilities with high prediction and reliability (Marek et al., 2022; Pohl et al., 2019; Sui et al., 2020). Here, prediction refers to a capacity to estimate the cognitive abilities of unseen individuals (outside of the model-building process, aka out-of-sample) (Yarkoni & Westfall, 2017). Reliability refers to the test-retest stability of measurements (Noble et al., 2021). This failure has led to headlines, such as ‘Scanning the Brain to Predict Behavior, a Daunting ‘Task’ for MRI’ (APS, 2020) and ‘Can brain scans reveal behaviour? Bombshell study says not yet: Most studies linking features in brain imaging to traits such as cognitive abilities are too small to be reliable, argues a controversial analysis’ (Callaway, 2022). Having highly predictive and reliable brain-based biomarkers for cognitive abilities could aid in our studies of mental-illness mechanisms (Morris & Cuthbert, 2012).

Predicting out-of-sample individual differences in cognitive abilities from neuroimaging has predominately been focused on non-task MRI modalities. Earlier studies associating general cognitive abilities and structural MRI (sMRI; reflecting brain volume/morphology) showed a weak association at *r* .1-.3 (McDaniel, 2005; Pietschnig et al., 2015). Because these associations were often done within-sample (not tested on unseen individuals), these weak associations may have already been biased upward (i.e., overfitting). More recent studies have shifted toward predictive modelling, a machine-learning/multivariate approach that draws information across different brain areas to maximize out-of-sample prediction. Yet, a recent predictive modelling competition showed *r* as low as .03 for out-of-sample prediction of general cognitive abilities via sMRI in children (Mihalik et al., 2019), suggesting the poor predictive performance of sMRI for cognitive abilities. Recently, researchers have turned to resting-state functional connectivity (resting-state FC) for prediction. Resting-state FC reflects the functional-connectivity between different brain areas, intrinsically occurring while resting. Applying predictive modelling on resting-state FC, researchers have found the moderate out-of-sample prediction of general cognitive abilities at *r*.2-.42 (Dubois et al., 2018; Rasero et al., 2021; Sripada et al., 2020, 2021). Still, there is a large room for improvement.

Here we examined two potential solutions: 1) using task-based functional MRI (tfMRI) and 2) combining tfMRI across tasks and with other modalities. First, tfMRI reflects the changes in BOLD induced by certain events while performing cognitive tasks. Sripada and colleagues (2020) applied predictive modelling on tfMRI using the Human Connectome Project (HCP) (Barch et al., 2013). They found that tfMRI from some tasks (e.g., tapping working memory, relational skills and language) predicted cognitive abilities very well, at out-of-sample *r*>.4 with the best task, working memory, at *r*=.5, which is higher than predictive performance found using the restingstate FC in this dataset at *r*=.26. This suggests that task-based activation during certain cognitive processes is a better candidate for capturing individual differences in cognitive abilities, compared to more commonly used modalities, e.g., sMRI and resting-state FC.

Nonetheless, tfMRI has recently come under intense scrutiny for its low test-retest reliability (Elliott et al., 2020). Researchers often quantify reliability using intraclass correlation (ICC) where low ICC reflects poor reliability (Cicchetti & Sparrow, 1981). Elliot and colleagues (2020) examined ICC of task-based activation at different regions and tasks using the HCP and showed poor ICC (<.4) across the regions and tasks. This is very different from sMRI’s ICC, which was in the excellent range (ICC>.75). Accordingly, while potentially providing better prediction, tfMRI may not be stable across time. Nonetheless, this ICC examination only focused on ICC from single regions, as opposed to from multiple regions drawn together by predictive modelling. Thus, this calls for research to examine the boosted reliability of tfMRI from the predictive modelling (Kragel et al., 2021).

The second solution involves combining tfMRI across tasks and with other MRI modalities. Most studies rely on a single tfMRI task or a single MRI modality to predict cognitive abilities (Sripada et al., 2020; Sui et al., 2020). Yet different tasks and MRI modalities might provide different sources of information for cognitive abilities, and thus combing them may help boost prediction and reliability (Rasero et al., 2021). There are different approaches for combining tfMRI across tasks and with other MRI modalities. The most straightforward approach is to include all MRI features from every task/modality in the same model. We call this approach, the “flat model.” Apart from the flat model, recent MRI researchers have used a machine-learning technique called “stacking” (Wolpert, 1992) to combine different MRI modalities into the “stacked model” (Engemann et al., 2020; Rasero et al., 2021). To apply stacking, researchers first build a model that predicts a target variable (e.g., cognitive abilities) from MRI features separately from each of the modalities, resulting in one predicted model from each modality. Using these modality-specific models, the researchers then compute their predicted values as surrogate features for each modality and build another model to combine these surrogate features across modalities. For instance, Rasero and colleagues (2021) used the HCP and combined many non-task MRI modalities (e.g., sMRI and resting-state FC) via stacking and showed enhanced predictive performance, compared to single-modality models. However, potentially partly due to not including tfMRI into their stacked model, they only found relatively modest performance from stacking, *R^2^*=.078, or roughly estimated *r*=.28. Accordingly, many questions arise. First, can integrating tfMRI across different tasks and/or with other modalities via stacking improve the prediction and reliability of the brain-based models for cognitive abilities? Second, which of the two combining approaches, flat vs. stacked models, performs better?

Apart from these two questions, it is still unclear the extent to which we should draw information across brain features and tasks/modalities. To model the relationship between brain features and cognitive abilities, many regression-based machine-learning algorithms are applicable. Common algorithms include those based on penalised regression (Kuhn and Johnson 2013), tree-based regression (Breiman et al. 2017) and kernel-based regression (Cortes and Vapnik 1995). Some algorithms, such as Ridge regression (Kuhn and Johnson 2013) and Elastic Net (Zou and Hastie 2005), are linear and additive while other algorithms, such as Random Forest (Breiman 2001), XGBoost (Chen and Guestrin 2016) and Support Vector Regression (SVR) with the Radial Basis Function (RBF) kernel (Cortes and Vapnik 1995; Drucker et al. 1996), are non-linear and interactive. It is still untested if using non-linear and interactive algorithms would allow for higher prediction and reliability. For stacked models, as compared to flat and modality-specific models, this issue is more complicated since different algorithms can be used for building modality-specific models vs. for stacking surrogate measures across modalities. In fact, different stacking studies have used different algorithms without testing their performance against other combinations of algorithms. For instance, Engemann and colleagues (2020) used Ridge regression for building their modality-specific models and Random Forest for stacking surrogate measures across modalities while Rasero and colleagues (2021) used Elastic Net for both building modality-specific models and stacking models across modalities. Thus, our study also aimed to find the combinations of algorithms that could lead to better performance for stacked models.

Beyond potentially enhancing prediction and reliability, integrating tfMRI across tasks via predictive modelling and stacking can also yield substantive insights into the neural basis of cognitive abilities. Examining the feature importance of the predictive model for each tfMRI task would allow us to demonstrate which of the brain features contribute highly to the prediction of general cognitive abilities for this particular task. Similarly, examining the feature importance of the stacked model that combined tfMRI across tasks would allow us to demonstrate which of the tasks contribute highly to the prediction. Altogether, we can achieve a predictive model from tfMRI with high prediction without losing sight of its feature importance. Accordingly, investigating highly contributing brain areas from highly contributing tasks would identify brain networks associated with general cognitive abilities across different cognitive domains (i.e., tfMRI tasks). In principle, this stacking feature-importance approach is similar to meta-analyses of tfMRI (Müller et al., 2018). When conducting a meta-analysis for cognitive abilities, researchers often examine the consistency in areas activated in association with individual differences in cognitive abilities across different tfMRI tasks (Basten et al., 2015; Jung & Haier, 2007; Santarnecchi et al., 2017). Santarnecchi and colleagues (2017), for instance, meta-analysed multiple tfMRI studies that used different tasks to predict individual differences in cognitive abilities. They found the areas in frontoparietal and dorsal attention networks to be consistently activated across these studies, in line with the parieto-frontal integration theory of intelligence (Jung & Haier, 2007). Accordingly, to compare the stacking feature-importance approach and the meta-analyses of tfMRI, we also examined the similarity of the feature importance from our tfMRI models with the meta-analysis by Santarnecchi and colleagues (2017).

To improve the prediction and reliability of MRI in capturing cognitive abilities, we integrated MRI signals across the whole brain from different tasks, structural MRI and resting-state functional connectivity. We directly compared predictive models comprising different sets of MRI modalities (e.g., all MRI modalities vs. tfMRI from all tasks vs. tfMRI from a subset of tasks that performed well vs. non-task MRI modalities). We expected to see high prediction and reliability from models that included tfMRI, especially when tfMRI was combined across tasks and with other modalities. We also tested different approaches for combining tfMRI information from different areas across tasks and with other MRI modalities: (1) using flat vs stacked models and (2) using different combinations of machine-learning algorithms. In addition to prediction and reliability, we designed our machine-learning pipeline to be interpretable, such that we could examine the contribution from each brain feature across different MRI modalities and compare our feature importance with those found in a recent meta-analysis of cognitive abilities (Santarnecchi et al., 2017).

## 2. Materials and Methods

### 2.1 Participants

We used the Human Connectome Project’s (HCP) S1200 release (Van Essen et al., 2013; WU-Minn Consortium Human Connectome Project, 2018). This release included multimodal-MRI and cognitive-performance data from 1,206 healthy participants (not diagnosed with psychiatric and neurological disorders). We discarded participants whose data were flagged as having quality control issues by the HCP (n=91): either having the “A” (anatomical anomalies) or “B” (segmentation and surface) flag or having any known major issues (Elam, 2021). We also removed participants who had missing values in any of the multimodal-MRI (n= 233) or cognitive-ability (n=9) variables. This left 873 participants (473 females, *M*=28.7 (*SD*=3.7) years old). They are from 414 families as many participants are from the same family. We provided participants’ ID in our GitHub repository (see below). Participants provided informed consent, including consent to share de-identified data. The Institutional Review Board at Washington University oversighted the HCP’s study procedure.

To examine the test-retest reliability of our predictive models, we also used the HCP Retest Dataset. This dataset included 45 participants who completed the HCP protocol for the second time (*M*=139.029 (SD=67.31) days apart). We had 34 participants whose data were complete across the two visits and were not flagged as having any quality control issues.

### 2.2 Features: Multimodal MRI

The HCP provided complete details of the scanning parameters and preprocessing pipeline elsewhere (Barch et al., 2013; Glasser et al., 2013; Van Essen et al., 2013). Here, we used MRI data with the MSMAll alignment (Glasser et al., 2016; Robinson et al., 2018) and with extensive processing (e.g., for task-based functional MRI, we obtained the general linear contrasts, see below). In total, the MRI data can be organized into 12 different modalities (i.e., sets of features):

#### Modalities 1-7: Task-based functional MRI (tfMRI) from seven different tasks

The HCP collected tfMRI from seven tasks (Barch et al., 2013), giving rise to seven sets of features in our model. The study scanned participants during each of the tasks twice with different phase encodings: right-to-left (RL) and left-to-right (LR). The HCP described preprocessing steps for tfMRI elsewhere (Glasser et al., 2013). Briefly, they included B_0_ distortion correction, motion correction, gradient unwrap, boundary-based co-registration to T_1_-weighted image, non-linear registration to MNI152 space, grand-mean intensity normalization and surface generation (see https://github.com/Washington-University/HCPpipelines). Here we focused on general-linear model contrasts of tfMRI (cope1.dtseries.nii). We parcellated tfMRI into 379 regions of interest (ROIs) using Glasser cortical atlas (360 ROIs) (Glasser et al., 2016) and Freesurfer subcortical segmentation (19 ROIs) (Fischl et al., 2002) and extracted the average value from each ROI. We treated general-linear model contrasts between standard experimental vs. control conditions for each tfMRI task as different modalities:

First, in the working memory task, we used the 2-back vs. 0-back contrast. Here, participants had to indicate whether the stimulus currently shown was the same as the stimulus shown two trials prior [2-back] or as the target stimulus shown in the beginning of that block [0-back]. Second, in the language task, we used the story vs. math contrast. Here, participants responded to questions about Aesop’s fables [story] or math problems [math]. Third, in the relational processing task, we used the relational vs. match contrast. Here participants reported if two pairs of objects differed in the same dimension [relational] or matched with a given dimension [match]. Forth, in the motor task, we used the averaged movement vs. cue contrast. Here participants were prompted [cue] to subsequently execute a movement [movement] with their fingers, toes, and tongue. Fifth, in the emotion processing task, we used the face vs. shape contrast. Here participants decided which two of the bottom objects matched the top object, and all objects in each trial can either be (emotional) faces [face] or shapes [shape]. Sixth, in the social cognition task, we used the theory of mind vs. random contrast. Here participants saw movie clips of objects interacting with each other either socially [theory of mind] or randomly [random]. Seventh, in the gambling task, we used the reward vs. punishment contrast. Here, participants had to guess if a number was higher or lower than 5, and the correct guess was associated with winning (vs. losing) money. They mostly won in certain blocks [reward] and mostly lost in others [punishment].

#### Modalities 8: Resting-state functional connectivity (resting-state FC)

The HCP collected resting-state FC from four 15-min runs, resulting in one-hour-long data (Glasser et al., 2013; Smith et al., 2013). Half of the runs were right-to-left phase encoding, and the other half were left-to-right phase encoding. The HCP applied a preprocessing pipeline to resting-state FC that is similar to the pipeline the study applied to tfMRI (Glasser et al., 2013) (see https://github.com/Washington-University/HCPpipelines). The HCP denoised resting-state FC data using ICA-FIX (Glasser et al., 2016). We parcellated the denoised resting-state FC data into 379 ROIs using the same atlases as the tfMRI (Fischl et al., 2002; Glasser et al., 2016). After the parcellation, we extracted each ROI’s time series from each of the four runs and concatenated them into one. We then computed Pearson’s correlation between concatenated time series of each ROI pair, resulting in a table of 71,631 non-overlapping resting-state FC indices. Thereafter, we applied *r-to-z* transformation to the whole table. To reduce the number of features in the model, we applied a principal component analysis (PCA) of 75 components to this table (Rasero et al., 2021; Sripada et al., 2019, 2020). More specifically, to prevent data leakage between training and test sets (see the predictive modelling pipeline below), we extracted principal components (PCs) from the resting-state FC table from each training set and applied this PCA definition to the associated test set of the same cross-validation fold.

#### Modalities 9-12: Structural MRI (sMRI)

The HCP provided the preprocessing pipeline for sMRI elsewhere (Glasser et al., 2013). Please see the preprocessing scripts here https://github.com/Washington-University/HCPpipelines. We separated sMRI data into four different modalities: cortical thickness, cortical surface area, subcortical volume and total brain volume. For cortical thickness and cortical surface area, we used Destrieux parcellation (148 ROIs) from FreeSurfer’s aparc.stats file (Destrieux et al., 2010; Fischl, 2012). As for subcortical volume, we used subcortical segmentation (19 gray matter ROIs) from FreeSurfer’s aseg.stats file (Fischl et al., 2002). As for total brain volume, we included five features calculated by FreeSurfer: estimated intra-cranial volume (FS_IntraCranial_Vol), total cortical gray matter volume (FS_TotCort_GM_Vol), total cortical white matter volume (FS_Tot_WM_Vol), total subcortical gray matter volume (FS_SubCort_GM_Vol) and ratio of brain segmentation volume to estimated total intracranial volume (FS_BrainSegVol_eTIV_Ratio). Note, we separately modelled subcortical volume and total brain volume even though both of them reflect the volume of the brain. This is due to the convention in FreeSurfer (Fischl, 2012): FreeSurfer only includes calculations from subcortical areas in the subcortical volume indices but includes calculations from both cortical and subcortical areas in the total-brain-volume indices. See Supplementary materials for the model that included both subcortical volume and total brain volume. Briefly, the predictive performance of the model that combined subcortical volume and total brain volume together was similar to that of the model with the total brain volume by itself.

### 2.3 Target: General Cognitive Abilities

We trained our models to predict general cognitive abilities, reflected by the average score of cognition assessments in the NIH Toolbox (Weintraub et al., 2014), as provided by the HCP (CogTotalComp_Unadj). The assessments included picture sequence memory, Flanker, list sorting, dimensional change card sort, pattern comparison, reading tests and picture vocabulary. Note we used the age-unadjusted average score since we controlled for age in the models themselves (see below).

### 2.4 Predictive Modeling Pipeline: Modality-Specific, Stacked and Flat models

For our predictive modelling pipeline (Figure 1), we used eight-fold nested cross-validation (CV) to build the models and evaluate their predictive performance. Since the HCP recruited many participants from the same family (Van Essen et al., 2013; WU-Minn Consortium Human Connectome Project, 2018), we first controlled the influences of the family structure by splitting the data into eight folds based on families. In each of the folds, there were members of ^~^50 families (M=109.13, SD=.33 participants), prohibiting members of the same family to be in the same folds. Note we used eight folds, as opposed to ten folds, here so that there were over 100 observations in each fold.

**Figure 1.**
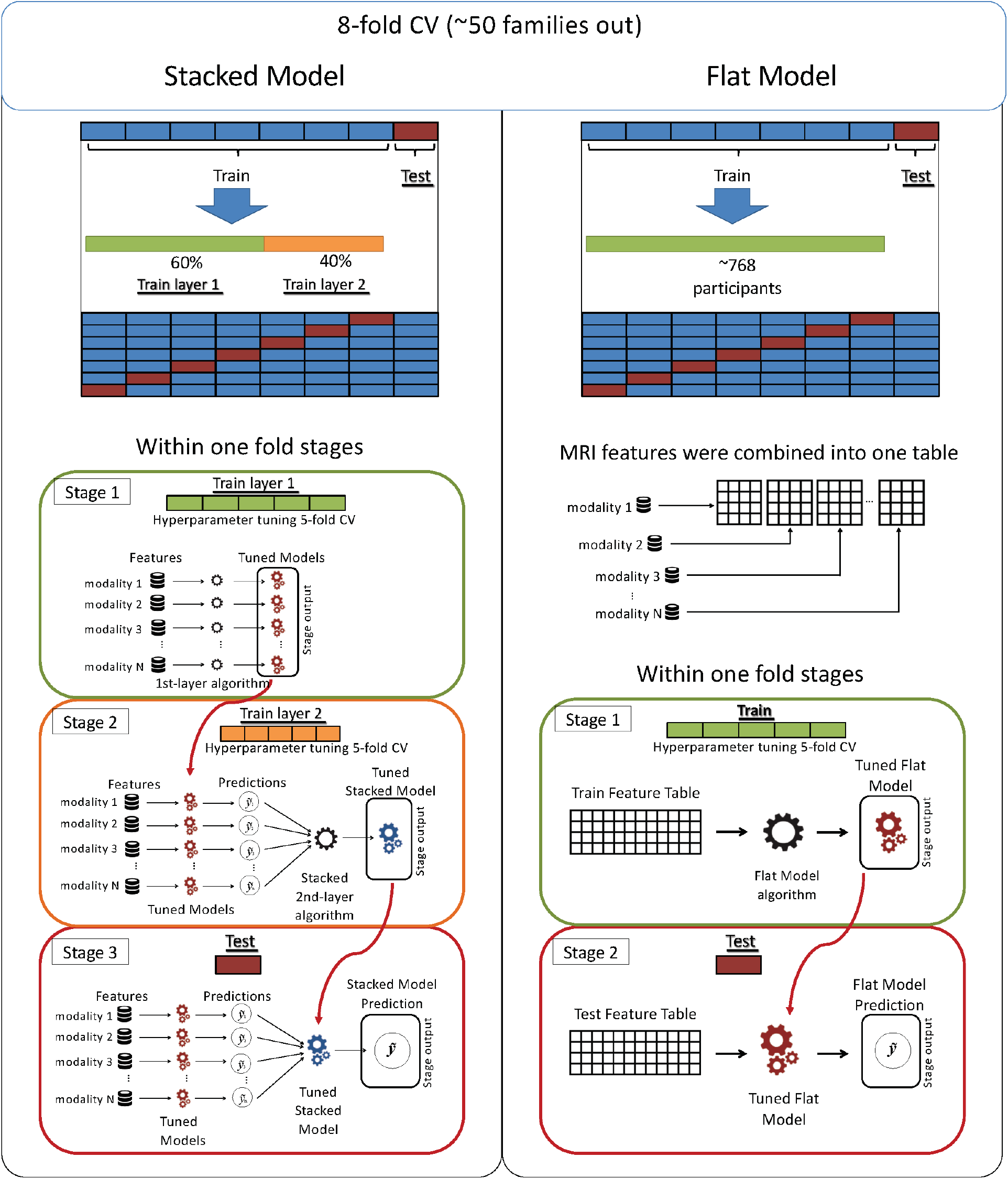
Predictive modelling pipeline. The diagram shows how we built stacked models and evaluated their predictive performance. CV = cross validation.

The nested CV involved two loops, nested with each other. In each CV “outer” loop, one of the eight folds that included ^~^50 families was held-out and was treated as a test set. The rest was treated as a training set. For stacked models, the training set was further split into 60% first- and 40% second-training layers. Within the CV “inner” loops, we separately fit the first-layer data from each modality to predict general cognitive abilities. Here we applied a five-fold CV to tune the hyperparameters of the models. This stage allowed us to create 12 of the modality-specific models. Using the second-layer data, we then computed predicted values for each of the 12 modalities based on the modality-specific models, and fit these predicted values across modalities to predict general cognitive abilities, resulting in the stacked models. Same as the first-layer data, we also applied a five-fold CV to tune the hyperparameters of the models here. For the flat models, we did not split the training set into two layers. Instead, we included features from different modalities directly in the model. Same with the stacked models, we applied a five-fold CV to tune the hyperparameters of the flat models.

For both stacked and flat models, we created four types of models that combined MRI of different modalities: 1) all-modality (i.e., a combination of 12 modalities), 2) all-task (i.e., a combination of seven different tfMRI tasks), 3) top-task (i.e., a combination of tfMRI tasks that showed high predictive performance) and 4) non-task (i.e., a combination of resting-state FC and four sMRI modalities).

### 2.5 Confound Correction and Standardisation

We first controlled for age (Dosenbach et al., 2010; Geerligs et al., 2015) and sex (Ruigrok et al., 2014; Trabzuni et al., 2013) in our models by linearly residualising them from both MRI data and cognitive abilities. We additionally residualised in-scanner movements from tfMRI and resting-state FC, given their sensitivity to motion artifacts (Power et al., 2012; Satterthwaite et al., 2013). More specifically, we defined in-scanner movements as the average of relative displacement (Movement_RelativeRMS_mean) across all available runs for each modality separately. We also standardised MRI data. To prevent data leakage, we separately applied residualisation and standardisation to the training and test sets. For the training sets, we implemented them separately on the two layers for the stacked models and as a whole training set for the flat models.

In our main analysis, we did not control for race/ethnicity due to the unbalanced number of participants from each race/ethnicity (see Supplementary Table 1). First, the HCP only included a few participants from certain races/ethnicities. For instance, only 2 people identified themselves as American Indian or Alaskan Native and not Hispanic Latino. In fact, there were 10 races/ethnicities (out of 14) that had fewer than 20 participants. Second, the majority (n=606) of participants who passed our exclusion criteria (n=873) were of the same race/ethnicity, “White and not Hispanic/Latino”. Accordingly, controlling race/ethnicity using linear residualisation from both MRI data and cognitive abilities can be problematic.

Nonetheless, to ensure that our results were robust against variation in race/ethnicity, we also conducted supplementary analysis by first excluding participants who belonged to a group of race/ethnicity with fewer than 20 participants. This left 834 participants but still resulted in a relatively low number of participants from certain race/ethnicity: 54 participants for Asians, Native Hawaiian other Other Pacific Islanders and not Hispanic/Latino and 57 for White and Hispanic/Latino”. We then residualised race/ethnicity in addition to age, sex and motion (see Supplementary Materials).

### 2.6 Predictive Modeling Algorithms

Here we employed four model-fitting algorithms via the scikit-learn package (Pedregosa et al., 2011): Elastic Net (Zou & Hastie, 2005), Random Forest (Breiman, 2001), XGBoost (Chen & Guestrin, 2016) and Support Vector Regression (SVR) (Cortes & Vapnik, 1995; Drucker et al., 1996). Given that the stacked models imposed two training layers, we had 16 (i.e., four by four) combinations of algorithms for our stacked models.

#### 2.6.1 Elastic Net

Elastic Net (Zou & Hastie, 2005) is a linear and additive algorithm that was previously used for predicting cognitive abilities from resting-state FC (Dubois et al., 2018). Elastic Net is a general form of penalized regression, allowing us to simultaneously draw information across different brain indices to predict one target variable. Compared to the classical, ordinary least squares regression, Elastic Net allows us to have more parameters (e.g., number of brain indices) than the number of observations (e.g., participants in each training set). Compared to other more complicated algorithms, Elastic Net has the benefit of being easier to interpret (Molnar, 2019). Researchers can directly interpret the magnitude of each coefficient as the importance of each feature (e.g., brain indices).

Elastic Net fits a plane that minimises the squared distance between itself and the data points (James et al., 2021; Kuhn & Johnson, 2013). When strongly correlated features are present, the classical, ordinary least squares regression tends to give very unstable estimates of coefficients and extremely high estimates of model uncertainty (Alin, 2010; Graham, 2003; Monti, 2011; P. Vatcheva & Lee, 2016). To address this, Elastic Net simultaneously minimises the weighted sum of the features’ slopes. For example, if the features are tfMRI from different regions, Elastic Net will shrink the contribution of some regions closer towards zero. The degree of penalty to the sum of the feature’s slopes is determined by a shrinkage hyperparameter ‘α’: the greater the α, the more the slopes shrunk, and the more regularised the model becomes. Elastic Net also includes another hyperparameter, ‘*ℓ*_1_ ratio’, which determines the degree to which the sum of either the squared (known as ‘Ridge’; *ℓ*_1_ ratio=0) or absolute (known as ‘Lasso’; *ℓ*_1_ ratio=1) slopes is penalised (Zou & Hastie, 2005). The objective function of Elastic Net as implemented by sklearn is defined as:

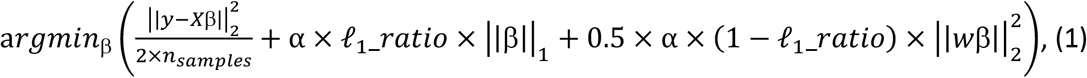

where *X* is the features, *y* is the target, and β is the coefficient.

To find the appropriate hyperparameters for each training layer, we applied a grid-search, 5-fold cross-validation (Efron & Gong, 1983; Hawkins et al., 2003; Koul et al., 2018) separately on each layer. In our grid, we searched for α using 500 numbers in log space, ranging from 10^−6^ and 10^4^, whereas for the **ℓ**_1_ ratio we used 100 numbers in linear space, ranging from 0 and 1.

#### 2.6.2 Random Forest

Random Forest (Breiman, 2001) is a tree-based algorithm. Unlike Elastic Net, Random Forest allows for a non-linear relationship between each feature and the target and for interactions among features. Random Forest bootstraps observations and incorporates a random subset of features at each split of tree building, resulting in several, bootstrapped trees. A prediction then is based on an aggregation of predicted values across bootstrapped trees. Here we used 5000 trees (i.e., setting ‘n_estimators’=5000 in sklearn). We also tuned two hyperparameters: the maximum depth of each tree, i.e., ‘max_depth’, and the number of features that are randomly sampled at each split, i.e., ‘max_feature’. In our grid, we used the integers between 1 to 11 for ‘max_depth’. For ‘max_feature’, we included the number of features itself, the square root of the number of features and log-based 2 of the number of features.

#### 2.6.3 XGBoost

XGBoost (Chen & Guestrin, 2016) is another tree-based algorithm. Like Random Forest, XGBoost allows for non-linearity and interaction. Unlike bootstrapped trees in Random Forest, XGBoost generates sequential trees where a current tree adapts from the gradients of previous trees. XGBoost has many parameters, most of which we used the default values, set by sklearn. We used ‘gbtree’ as a booster and tuned 3 hyperparameters: ‘max_depth’, ‘eta’ and ‘subsample’.

Same with Random Forest, ‘max_depth’ is the maximum depth of each tree. Here, we sampled ‘max_depth’ from the integers between 1 to 5. ‘eta’ is a learning rate that controls the speed of tree adaptation. We tuned ‘eta’ using the values: .03, .06 and .1. Lastly, ‘subsample’ is the ratio of the training instance. We tuned ‘subsample’ using the values: .6, .8 and 1.

#### 2.6.4 Support Vector Regression

Support Vector Regression (SVR) (Cortes & Vapnik, 1995; Drucker et al., 1996) is a kernel-based algorithm, which also allows for non-linearity and interaction. The objective function of SVR is defined as:

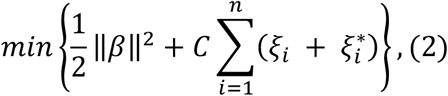

with constraints:

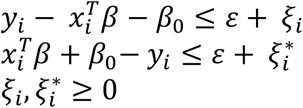

where *x* is the feature, *y* is the target, β is the coefficient, *ε* is the margin of tolerance where no penalty is given to errors, are non-zero slack variables that are allowed to be above (*ξ_i_*) and below (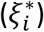) the margin of tolerance, and *C* or ‘complexity’ is a regularisation hyperparameter. A higher *C* indicates a stronger penalty for high complexity (i.e., less tolerant to wrong-side errors). Accordingly, regularisation is achieved by disfavouring high-complexity models that have fewer errors on training data but may pick up much larger errors from test data. Here we sampled *C* from following set: {1,6,9,10,12,15,20 and 50}. To allow for non-linearity and interaction, we applied a kernel trick with the Radial Basis Function (RBF) to transform the data into a higher-dimensional space, defined as

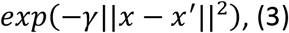

where *γ* is the kernel coefficient. We sampled the kernel coefficient from one divided by the multiplication of the number of features and its variance, one divided by the number of features as well as the following values: 10^−8^, 3×10^−8^, 6×10^−8^, 10^−7^, 3×10^−7^, 6×10^−7^, 10^−6^, 3×10^−6^, 6×10^−6^, 10^−5^, 3×10^−5^, 6×10^−5^, 10^−4^, 3×10^−4^, 6×10^−4^, 10^−3^, 3×10^−3^ and 6×10^−3^.

### 2.7 Modelling Performance: Prediction

To evaluate the models’ prediction, we used the eight held-out folds. We first computed predicted values from each model. We then tested how similar these predicted values were to the real, observed values, indicated by four predictive performance metrics: Pearson’s *r*, coefficient of determination (R^2^), mean square error (MSE) and mean absolute error (MAE) (See Supplementary materials for the exact definitions). Using multiple metrics of predictive performance is highly recommended, as each metric can reveal different aspects of the models’ performance (Poldrack et al., 2020).

To statistically compare measures of predictive performance across models, we first combined predicted and observed values across the eight held-out folds. We then created bootstrap distributions (Efron & Tibshirani, 1993) of the differences in the four measures of predictive performance between each pair of models. If these distributions of the differences did not include zero inside their 95% confidence interval, we concluded that the two models were significantly different from each other.

### 2.8 Modeling Performance: Test-retest reliability

To evaluate test-retest reliability, we applied a similar pipeline as done recently (Taxali et al., 2021). Here, as opposed to using the eight-fold nested CV, we implemented a train-test split. We treated the data from participants who were scanned twice as a test set (as opposed to using 50 held-out families) and the rest as a training set. Same as before, we further split the training set into 60% first- and 40%second-training layers. We used the first-training layer for building modality-specific models via a five-fold CV and the second-training layer for combining the predicted values from the first-training layer into stacked models via another five-fold CV. We then applied the final, tuned models to the test set. Given that each testing participant had data from two time points, we obtained two predicted values of cognitive abilities for each testing participant. This allowed us to test the extent to which the predicted values from different models were stable across the two time points, using the intraclass correlation (ICC) (Shrout & Fleiss, 1979).

ICC is generally defined as

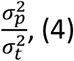

where 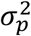 is the between-participant variance, and 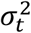 is the within-participant variant. There are two commonly used types of ICC for test-retest reliability in MRI (Noble et al., 2021).

First, ICC(2,1) reflects an absolute agreement with random sources of error. ICC(2,1) is defined as

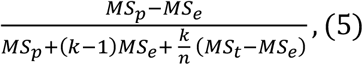

where *MS_p_* is mean square for participants, *MS_e_* is mean square for error, *MS_t_* is mean square for time points (i.e., measurements), n is the number of participants, k is the number of time points.

Second, ICC (3,1) reflects a consistency with fixed sources of error. ICC (3,1) is defined as

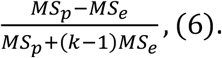

We computed both types of ICC using the Pingouin package (https://pingouin-stats.org/). Based on an established criterion (Cicchetti & Sparrow, 1981), we considered ICC < .4 as poor, ICC ≥ .4 and <.6 as fair, ICC ≥ .6 and < .75 as good and ICC ≥ .75 as excellent.

### 2.9 Feature Importance: Elastic Net Coefficients

To understand which brain features contributed stronger to the prediction of cognitive abilities, we used Elastic Net Coefficients. We chose to interpret Elastic Net as opposed to other algorithms due to a) its interpretability and b) its high predictive performance compared to other algorithms (see Results). Given that Elastic Net makes a prediction based on a weighted sum of features, its coefficients are readily interpretable: stronger magnitude of the coefficient means a higher contribution to the prediction.

We used the Elastic Net coefficients to locate 1) which of the modalities contributed highly to the prediction of the stacked models and 2) which of the brain features contributed highly to the prediction of the modality-specific models. More specifically, once we showed which of the 12 modalities contributed highly to the prediction of the all-modality stacked model, we would then investigate the brain indices of the top-performing modality-specific models that contributed highly to the prediction. In addition to plotting the coefficients on the brain images, we also provided a list of top-20 brain indices for each top-performing modality-specific model. We evaluated the contribution of each brain index based on the magnitude of their Elastic Net coefficient. In this list, we identified brain networks associated with each brain region using the Cole-Anticevic Brain Network Atlas (Glasser et al., 2016; Ji et al., 2019). We also provided MNI coordinates for each region, obtained by transforming voxel coordinates (based on https://neuroimaging-core-docs.readthedocs.io/en/latest/pages/atlases.html) to the MNI space via nilearn.image.coord_transform() using the standard FSL template, MNI152_T1_1mm, as a reference.

To represent contributing areas across tfMRI tasks, we combined the magnitude of Elastic Net coefficients from all tasks at each brain area, weighted by the overall magnitude of Elastic Net coefficients of the task stacked model. We also visualised how these contributing areas were overlapped with those found in a recent meta-analysis of cognitive abilities (Santarnecchi et al., 2017). Here we downloaded the Activation Likelihood Estimate (ALE) map of significant foci that showed associations with various cognitive abilities (Gf_net.nii from http://www.tmslab.org/netconlab-fluid.php) in MNI, volumetric space. We then converted this ALE map to the surface space and overlaid the ALE map on top of the magnitude of Elastic Net coefficients from all tasks using Connectome Workbench (Marcus et al., 2011).

### 2.10 Code Accessibility

The shell and Python scripts used in the analyses are made available here: https://github.com/HAM-lab-Otaqo-University/HCP

## 3. Results

### 3.1 Prediction for Modality-Specific Models

Figure 2 shows the predictive performance of modality-specific models based on the 8-fold CV. Among the 12 modality-specific models, working-memory tfMRI, language tfMRI, and relational tfMRI had the highest prediction, in descending order. Given their high predictive performance, we included tfMRI from these three tasks in the top-task stacked and flat models. On the contrary, the modality-specific models based on subcortical volume, cortical thickness and gambling tfMRI had a relatively poorer prediction. As for algorithms, Elastic Net and SVR demonstrated numerically better prediction than Random Forest and XGBoost.

**Figure 2.**
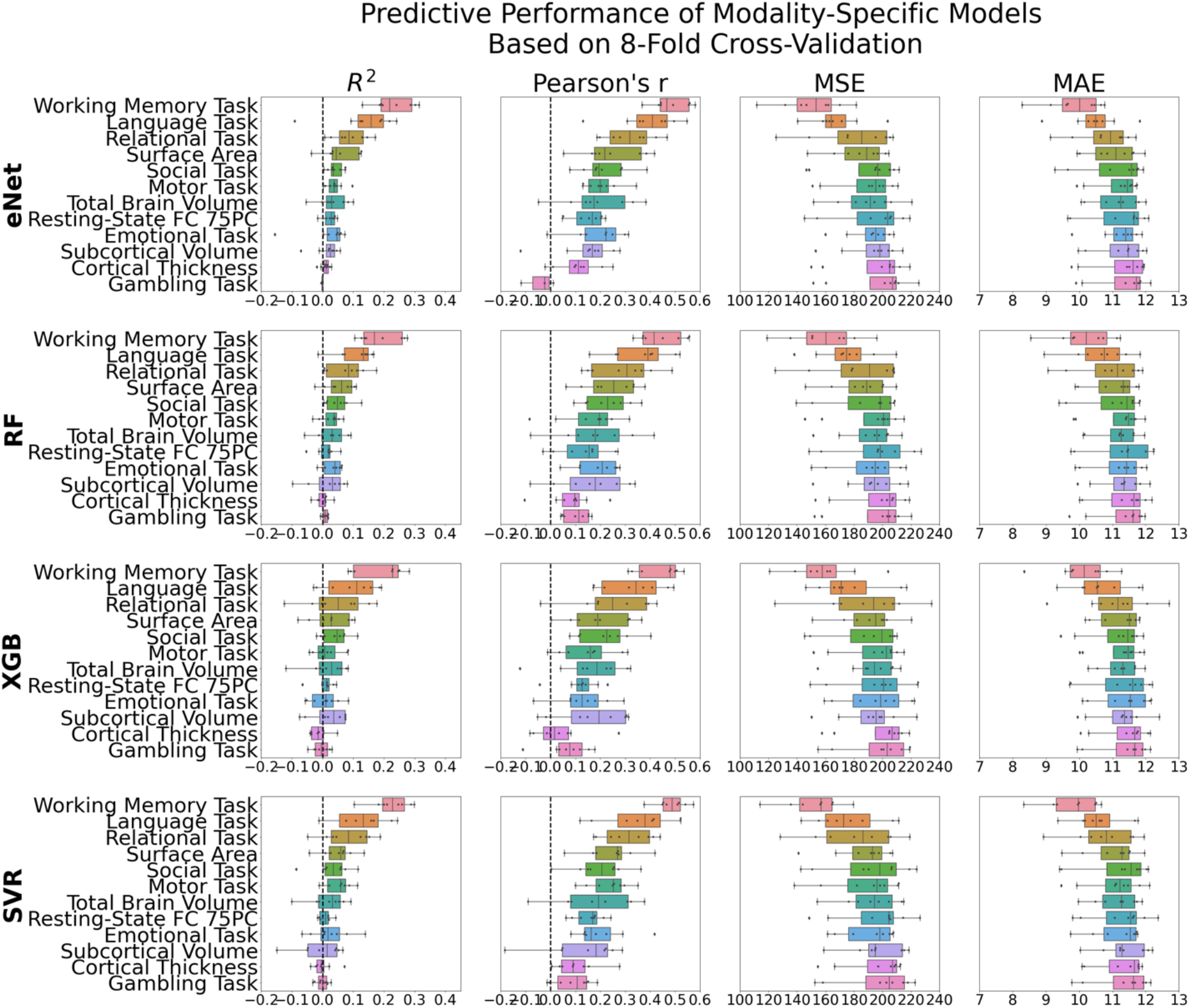
Predictive performance of modality-specific models based on the 8-fold CV across the four algorithms. Each dot represents predictive performance from each of the eight held-out folds. R^2^=coefficient of determination; eNet = Elastic Net; RF = Random Forest; XGB = XGBoost; SVR = Support Vector Regression.

### 3.2 Prediction for Stacked Models

Figure 3 shows the predictive performance of stacked models based on the 8-fold CV. Among the four stacked models, the all-modality stacked model had the numerically highest prediction, compared to other stacked models. Regarding algorithms, using Elastic Net on both training layers, denoted by eNet+eNet, for the all-modality stacked model provided numerically highest prediction.

**Figure 3.**
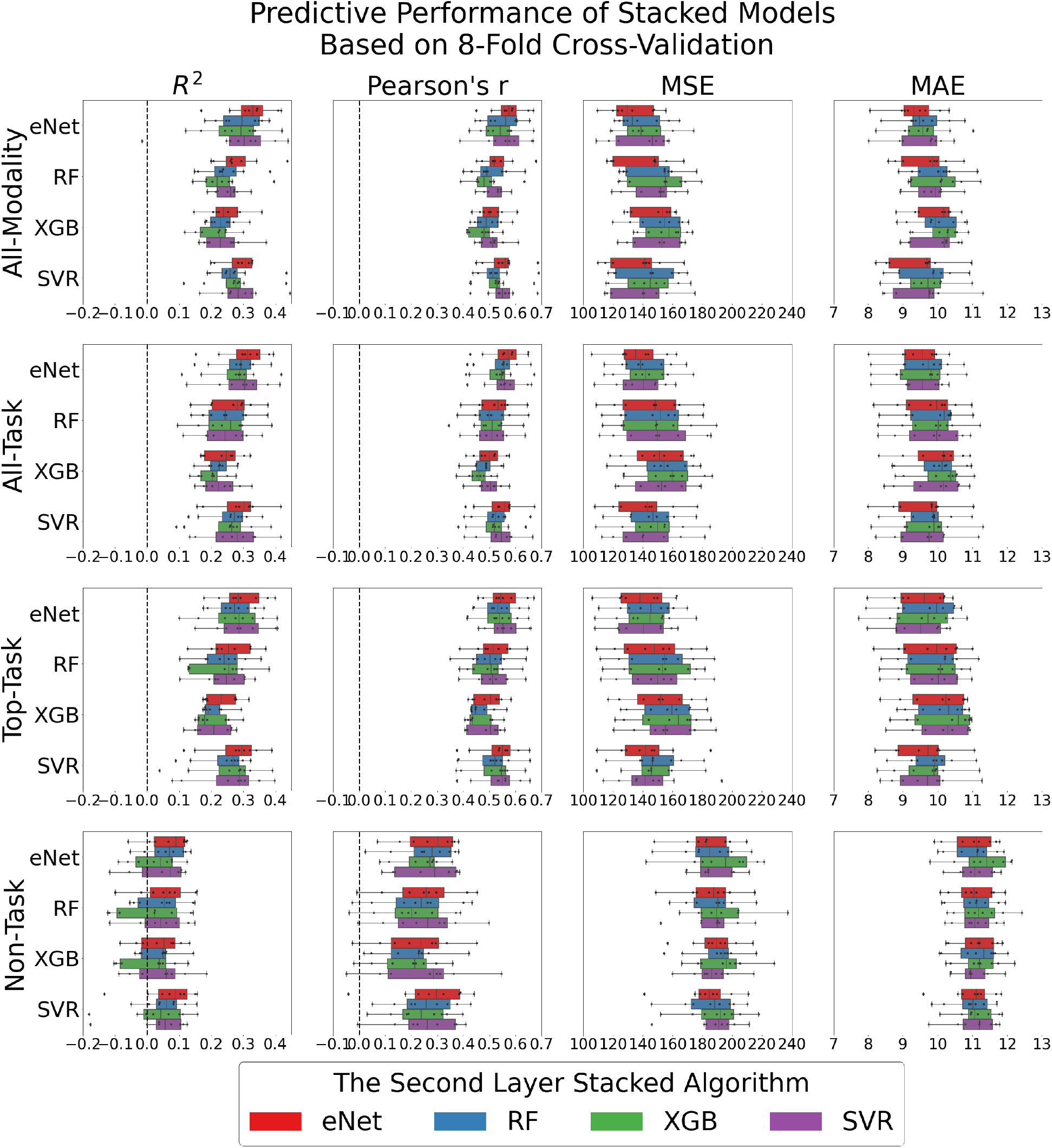
Predictive performance of stacked models based on the 8-fold CV across the four-by-four algorithms. Each dot represents predictive performance from each of the eight held-out folds. R^2^=coefficient of determination; eNet = Elastic Net; RF = Random Forest; XGB = XGBoost; SVR = Support Vector Regression.

Figure 4 shows scatter plots of the four stacked models that were trained on Elastic Net across the two training layers (eNet+eNet). Note that when race/ethnicity was added as another confounding variable, in addition to age, sex and in-scanner movements from tfMRI and resting-state FC, we found some reduction in predictive performance (see Supplementary Materials).

**Figure 4.**
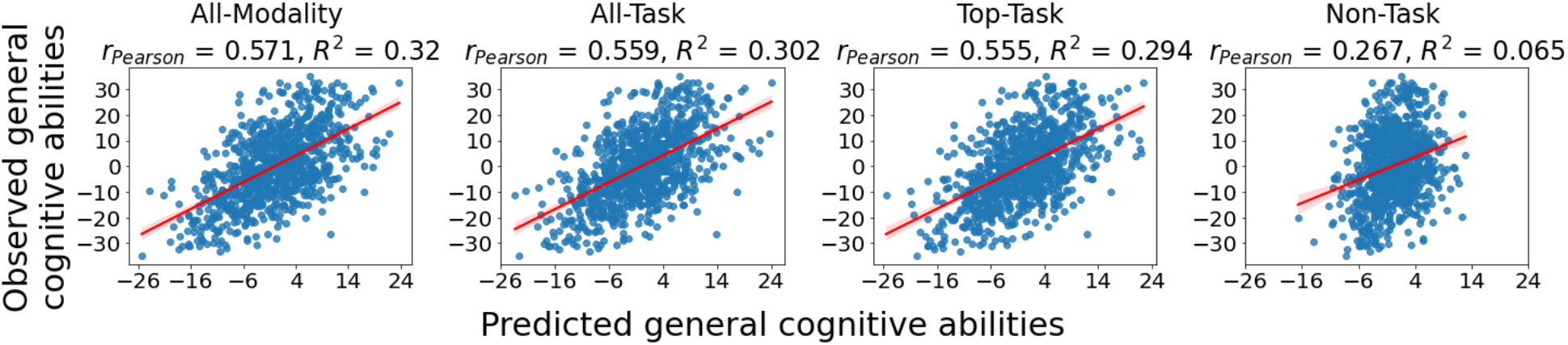
Scatter plots depicting the relationships between predicted and observed values of general cognitive abilities across the eight held-out folds of the stacked models trained on Elastic Net across both training layers. Note that Pearson’s r and R^2^ values are based on the average of the performance across 8-fold CVs.

This can be seen across eNet+eNet stacked model, including all-modality (before correction: Pearson’s r=.571 and R^2^=.32 and after correction: Pearson’s r=.522 and R^2^=.259), all-task (before correction: Pearson’s r=.559 and R^2^=.302 and after correction: Pearson’s r=.515 and R^2^=.248), top-task (before correction: Pearson’s r=.555 and R^2^=.294 and after correction: Pearson’s r=.512 and R^2^=.243) and non-task (before correction: Pearson’s r=.267 and R^2^=.065 and after correction: Pearson’s r=.138 and R^2^=.014).

Using bootstrapping, we compared the predictive performance of the eNet+eNet all-task stacked model with other eNet+eNet stacked models (Figure 5). Here, with the eNet+eNet algorithm, the all-modality stacked model predicted general cognitive abilities better than the all-task stacked model. And the all-task stacked model did not perform significantly better than the top-task stacked model but performed better than the non-task stacked model.

**Figure 5.**
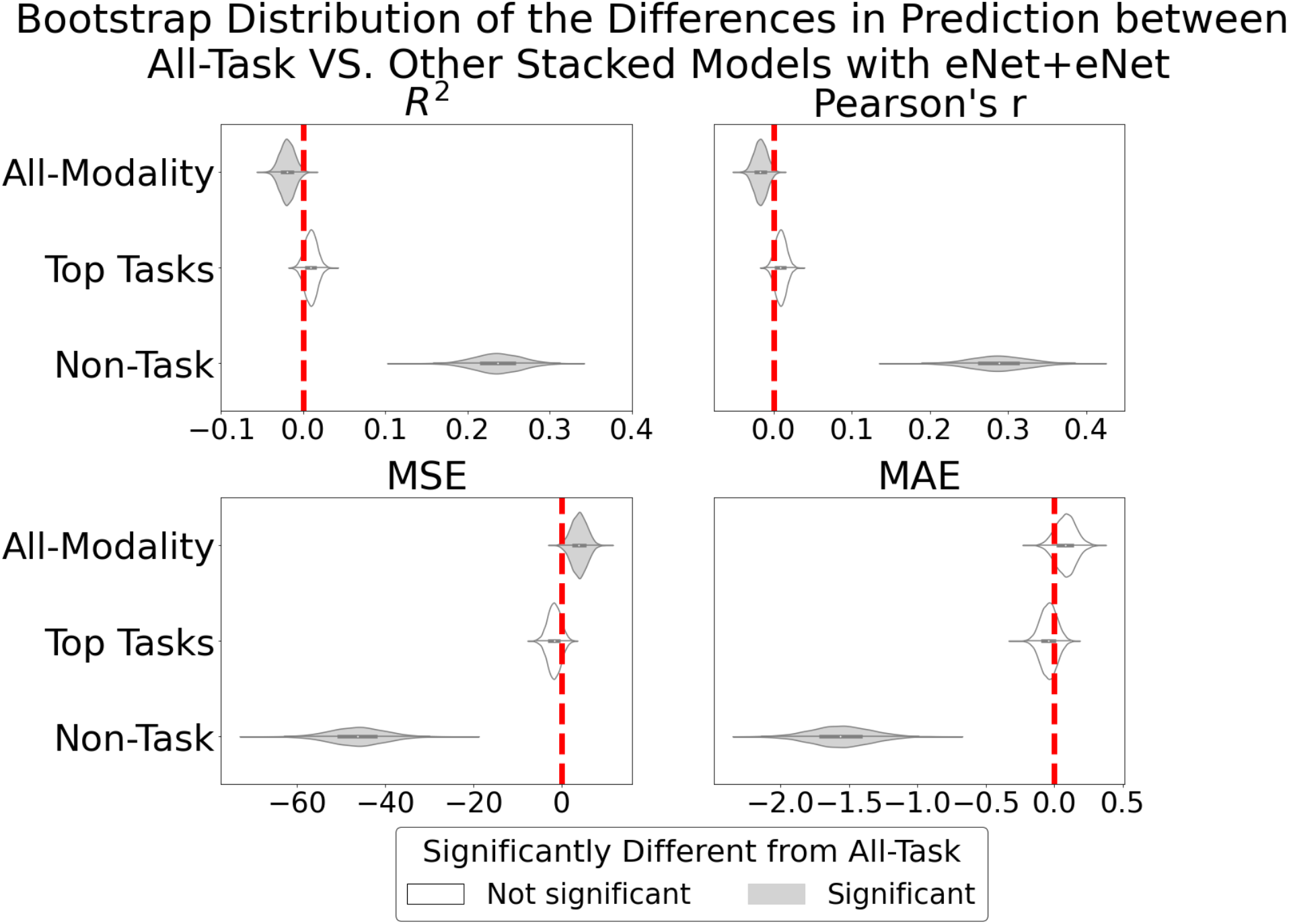
Bootstrap distribution of the differences in prediction between the all-task stacked model and other stacked models trained on Elastic Net across both training layers. For R^2^ and Pearson’s r, values lower than zero indicate better performance than the all-task stacked model. For mean square error (MSE) and mean absolute error (MAE), values lower than zero indicate worse performance than the all-task stacked model.

To investigate the predictive performance of stacking in comparison to using a single modality, we implemented bootstrapping to compare the performance of the eNet+eNet all-modality stacked model against 12 modality-specific models trained on the same algorithm, Elastic Net (Figure 6). The all-modality stacked model performed better than any of the modality-specific models.

**Figure 6.**
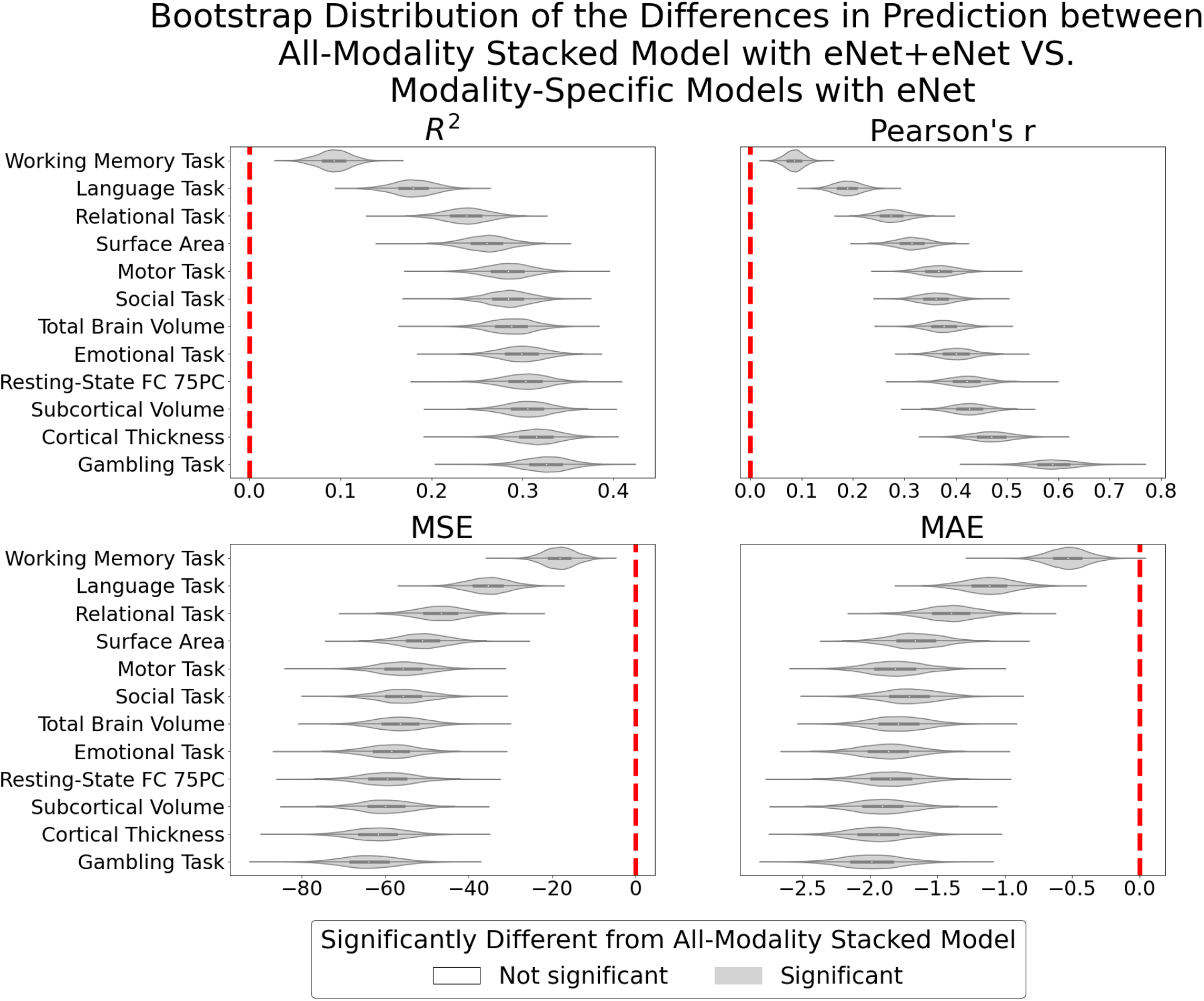
Bootstrap distribution of the differences in prediction between the all-modality stacked model trained on Elastic Net across both training layers and modality-specific models trained on Elastic Net. For R^2^ and Pearson’s r, values lower than zero indicate better performance than the all-modality stacked model. For mean square error (MSE) and mean absolute error (MAE), values lower than zero indicate worse performance than the all-modality stacked model.

Similarly, to investigate the predictive performance based on different algorithms, we implemented bootstrapping to compare the predictive performance of the eNet+eNet all-modality stacked model against the all-modality stacked model trained on other combinations of algorithms (Figure 7). Implementing Elastic Net across the two training layers led to significantly better predictive performance than any of other combinations of algorithms, apart from two: using SVR on the first training layer and Elastic Net on the second training layer and using SVR across the two training layers.

**Figure 7.**
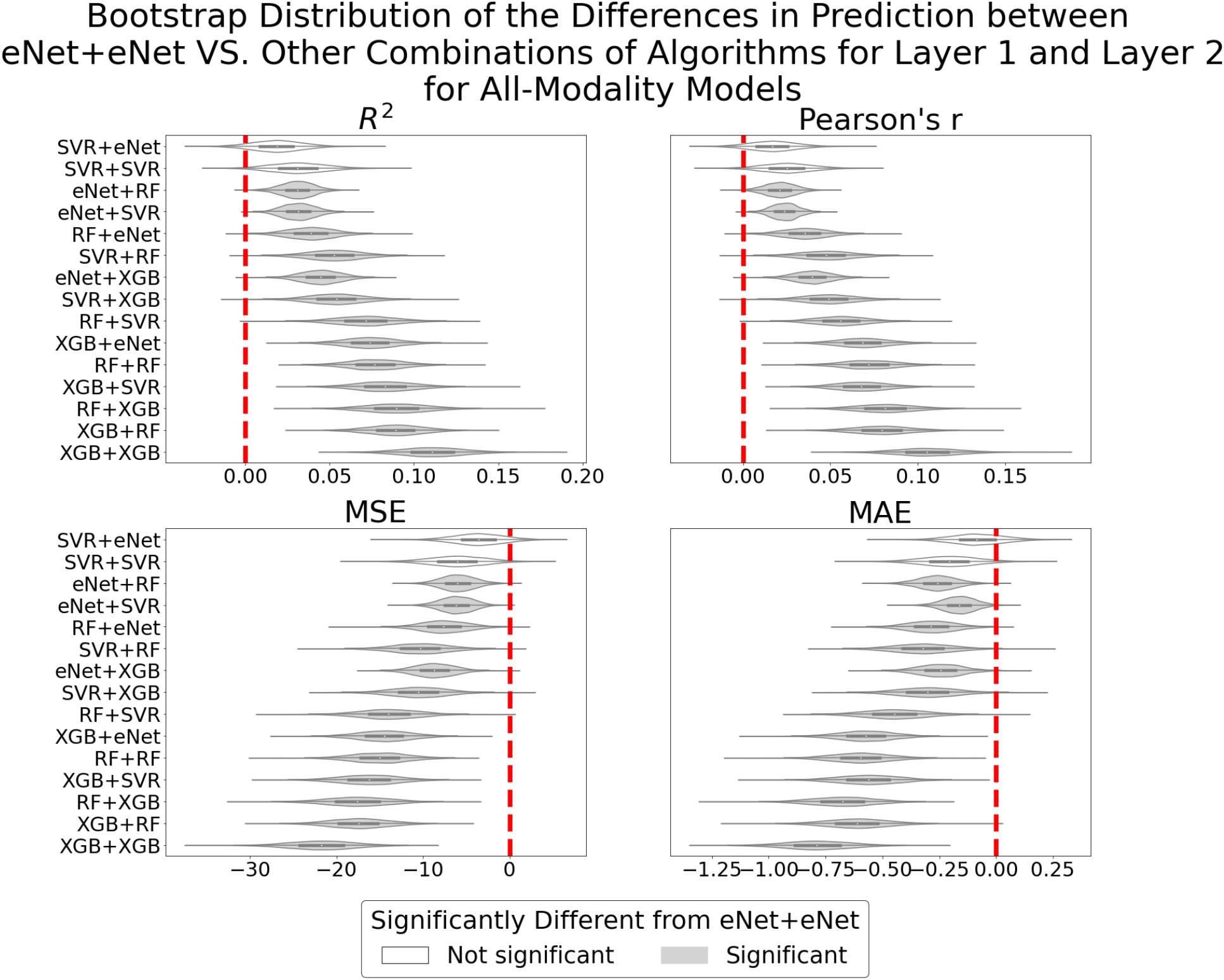
Bootstrap distribution of the differences in prediction between the all-modality stacked model trained on Elastic Net across both training layers and the all-modality stacked model trained on other combinations of algorithms. For Pearson’s r and R^2^, values lower than zero indicate better performance than the all-modality stacked model with eNet+eNet. For mean square error (MSE) and mean absolute error (MAE), values lower than zero indicate worse performance than the all-modality stacked model with eNet+eNet. The algorithm on the left of the plus symbol denotes the algorithm used for the first training layer, and the algorithm on the right of the plus symbol denotes the algorithm used for the second training layer. eNet = Elastic Net; RF = Random Forest; XGB = XGBoost; SVR = Support Vector Regression.

### 3.3 Comparison between Stacked and Flat Models in Prediction

To compare the stacked models with the flat models, we created bootstrap distribution of their predictive performance across algorithms and combinations of modalities (Figure 8). Overall, the performance of the stacked and flat models was similar to each other across algorithms and combinations of modalities.

**Figure 8.**
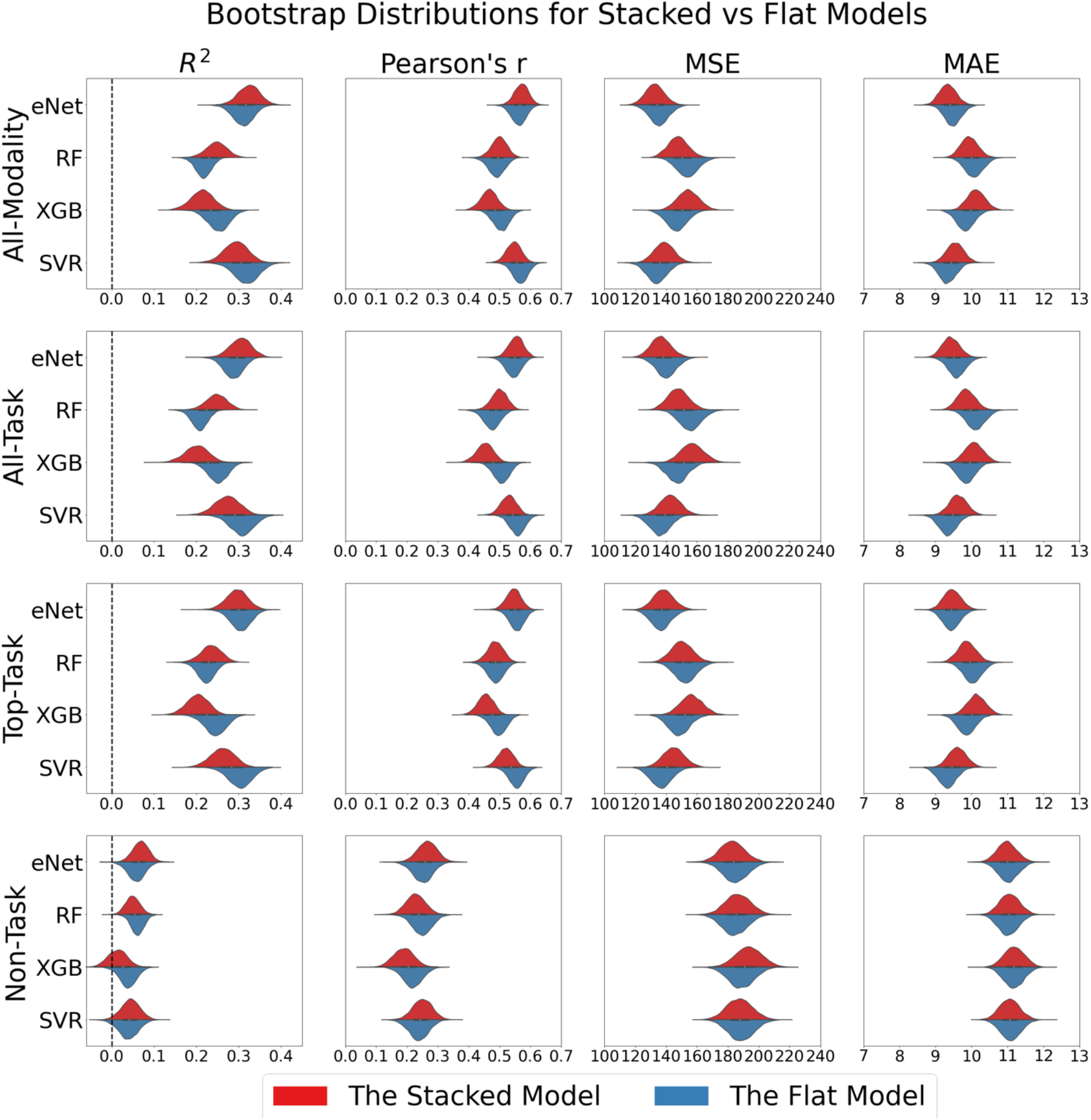
Bootstrap distribution of the predictive performance of stacked and flat models across algorithms and combinations of modalities. For the stacked models, we only plotted bootstrap distributions of models with the same machine-learning algorithms across the two training layers. eNet = Elastic Net; RF = Random Forest; XGB = XGBoost; SVR = Support Vector Regression.

### 3.4 Feature Importance of Stacked Models

Examining the all-modality stacked model’s Elastic Net coefficients (Figure 9) reveals the working memory tfMRI to be the main contributing modality, followed by language tfMRI, relational tfMRI, emotional tfMRI, surface area, resting-state FC, motor tfMRI and total brain volume, respectively. Cortical thickness, subcortical volume, social tfMRI and gambling tfMRI had relatively weaker contributions. We also saw a similar order of contributions from different modalities of other stacked models.

**Figure 9.**
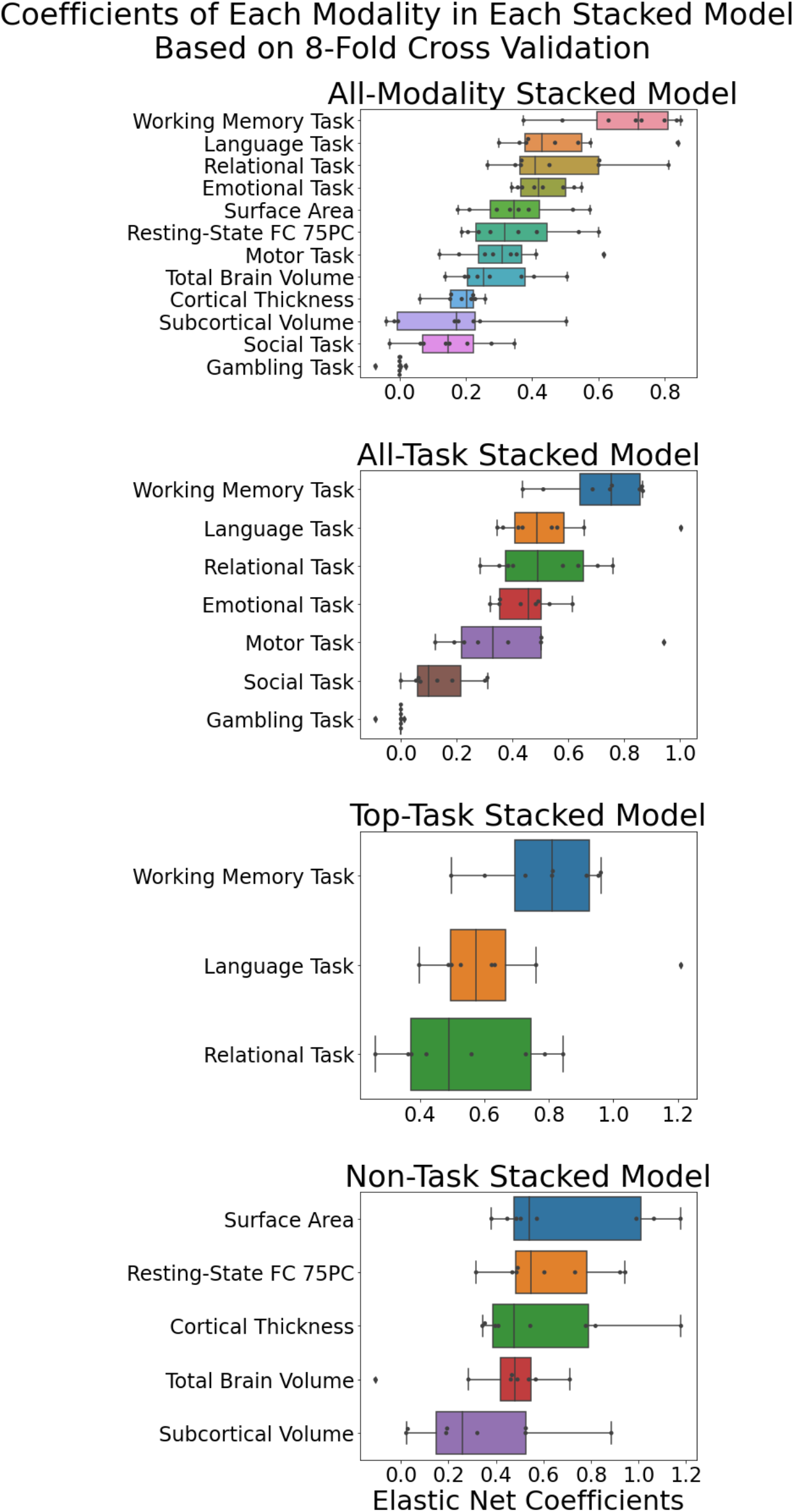
Feature importance of each stacked model, as reflected by Elastic Net coefficients.

### 3.5 Test-retest Reliability: the intraclass correlation (ICC)

Figure 10 shows the ICC of the stacked and single-modality models across the two definitions ICC(2,1) and ICC(3,1). Here we plotted ICC with algorithms that demonstrated relatively good predictive performance: eNet+eNet, SVR+eNet and SVR+SVR for stacked models and Elastic Net and SVR for the modality-specific models. Modality-specific models with sMRI-based modalities (total brain volume, surface area, subcortical volume and cortical thickness) had the highest ICC (> .88). Similarly, the all-modality, all-task and top-task stacked models with eNet+eNet had high test-retest reliability, reflected by the excellent level of ICC (> .75). The non-task stacked model across algorithms had a good level of ICC. Modality-specific tfMRI models with Elastic Net had ICC varied from unable-to-compute due to the models resulting in the same predicted value (i.e., mean) for every participant (gambling), poor (motor), fair (social, emotional and relational) to good (working memory and language). Modality-specific tfMRI models with SVR had a similar, somewhat poorer, level of ICC. The resting-state FC with Elastic Net and SVR had a fair and poor level of ICC, respectively. Figure 11 shows the predicted values of different models at the two sessions for each participant. Here models with high ICC showed consistency in predictive values across the two sessions, compared to models with lower ICC

**Figure 10.**
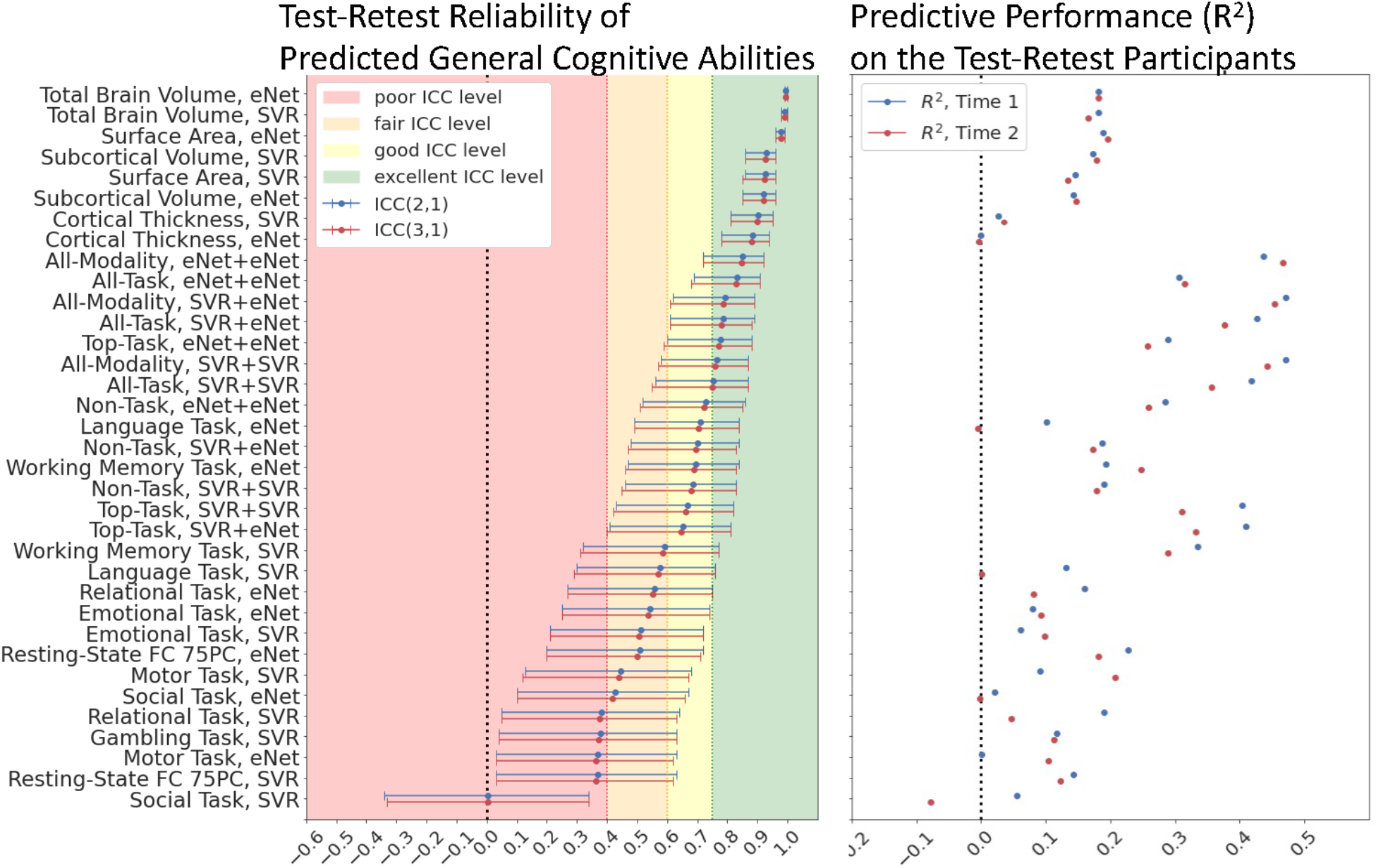
Test-retest reliability of stacked and modality-specific models. We computed test-retest reliability using two definitions of interclass correlation: ICC(2,1) and ICC(3,1). We only plotted ICC with algorithms that demonstrated relatively good predictive performance: eNet+eNet, SVR+eNet and SVR+SVR for stacked models and Elastic Net and SVR for the modality-specific models. Note the ICC of the modality-specific Gambling Task for Elastic Net cannot be computed. We also computed the predictive performance (R^2^) of the test-retest participants associated with each of the stacked and modality-specific models, separately for each time point.

**Figure 11.**
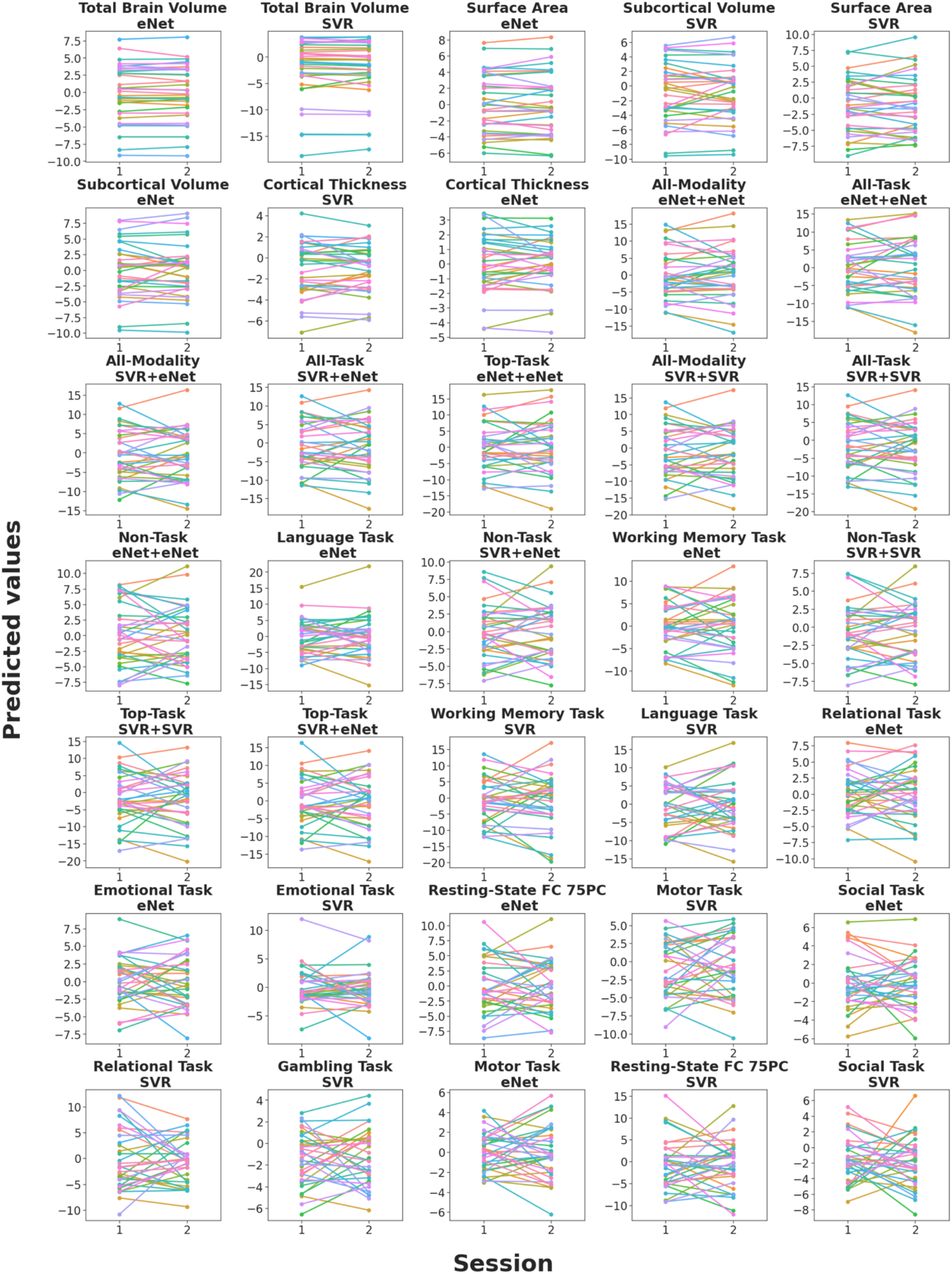
Predicted values of stacked and modality-specific models across two time points (i.e., session), ranked by ICC. Each line represents each participant. We only plotted predicted values of algorithms that demonstrated relatively good predictive performance: eNet+eNet, SVR+eNet and SVR+SVR for stacked models and Elastic Net and SVR for the single-modality models.

### 3.6 Feature Importance for Modality-Specific and All-Task Stacked Models

Figure 12 shows the feature importance of each modality-specific model, as reflected by Elastic Net coefficients, averaged across the eight cross-validation folds. Supplementary Tables 3 to 5 list 20 features (brain regions/connectivity pair) with the highest Elastic Net magnitude for each of the top-three modalities. We provided a full list of feature importance for all modalities in our GitHub repository.

**Figure 12.**
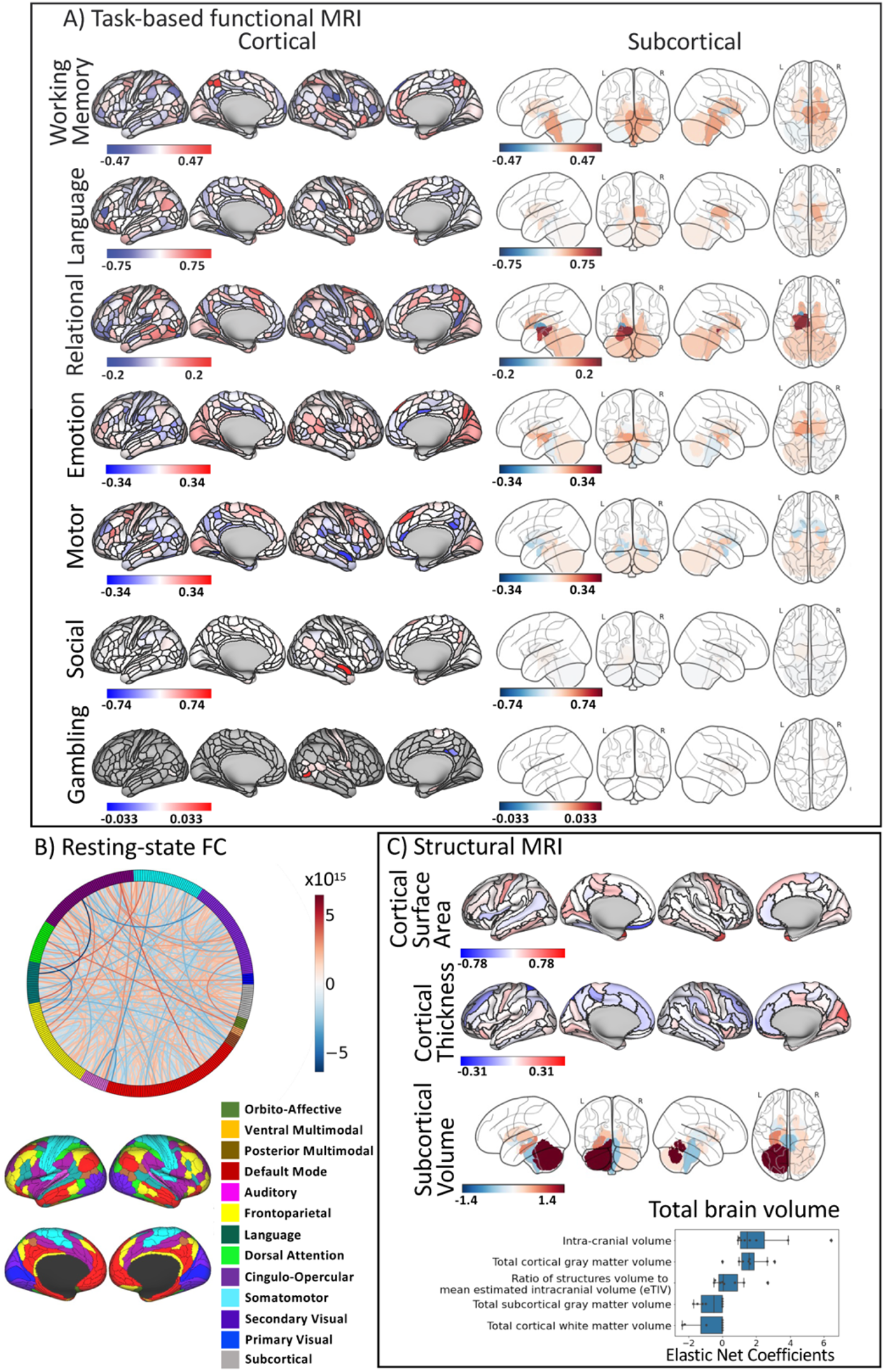
Feature importance of each modality-specific model, reflected by Elastic Net coefficients average across the eight cross-validation folds. a, b and c show Elastic Net coefficients for tfMRI, resting-state FC and sMRI, respectively. For resting-state FC, we plotted PCA loading for each pair of regions, weighted by Elastic Net magnitude, summed across the eight folds.

For working-memory tfMRI (Figure 12, Supplementary Table 3), we found highly contributing areas from the frontoparietal, default-mode, visual 2 and cingulo-opercular networks. These included areas such as the anterior cingulate, medial prefrontal, superior parietal, inferior frontal, and dorsolateral prefrontal cortices.

For language tfMRI (Figure 12, Supplementary Table 4), we found highly contributing areas from frontoparietal and default-mode networks. These included areas such as anterior cingulate, medial prefrontal, insular, orbital and polar frontal, inferior frontal, lateral temporal and dorsolateral prefrontal cortices.

For relational tfMRI (Figure 12, Supplementary Table 5), we found highly contributing areas from many networks, e.g., default-mode, visual 2, language, subcortical, frontoparietal and dorsal attention. These included areas such as orbital and polar frontal, posterior cingulate, visual, inferior frontal, superior parietal and dorsolateral prefrontal cortices.

Figure 13 and Table 1 show contributing brain regions across tfMRI tasks, reflected by the magnitude of Elastic Net coefficients from all tasks at each brain area, weighted by the magnitude of Elastic Net coefficients of the all-task stacked model with eNet+eNet. This figure shows the contribution of the areas in the default, frontoparietal and cingulo-opercular networks to the prediction of general cognitive abilities across tfMRI tasks. Additionally, overlaying the ALE map from a previous meta-analysis of cognitive abilities (Santarnecchi et al., 2017) on top of the contributing brain regions across tasks shows the overlapping regions in the frontoparietal network, in areas such as the left inferior frontal, left anterior cingulate and medial prefrontal and left superior parietal cortices.

**Figure 13.**
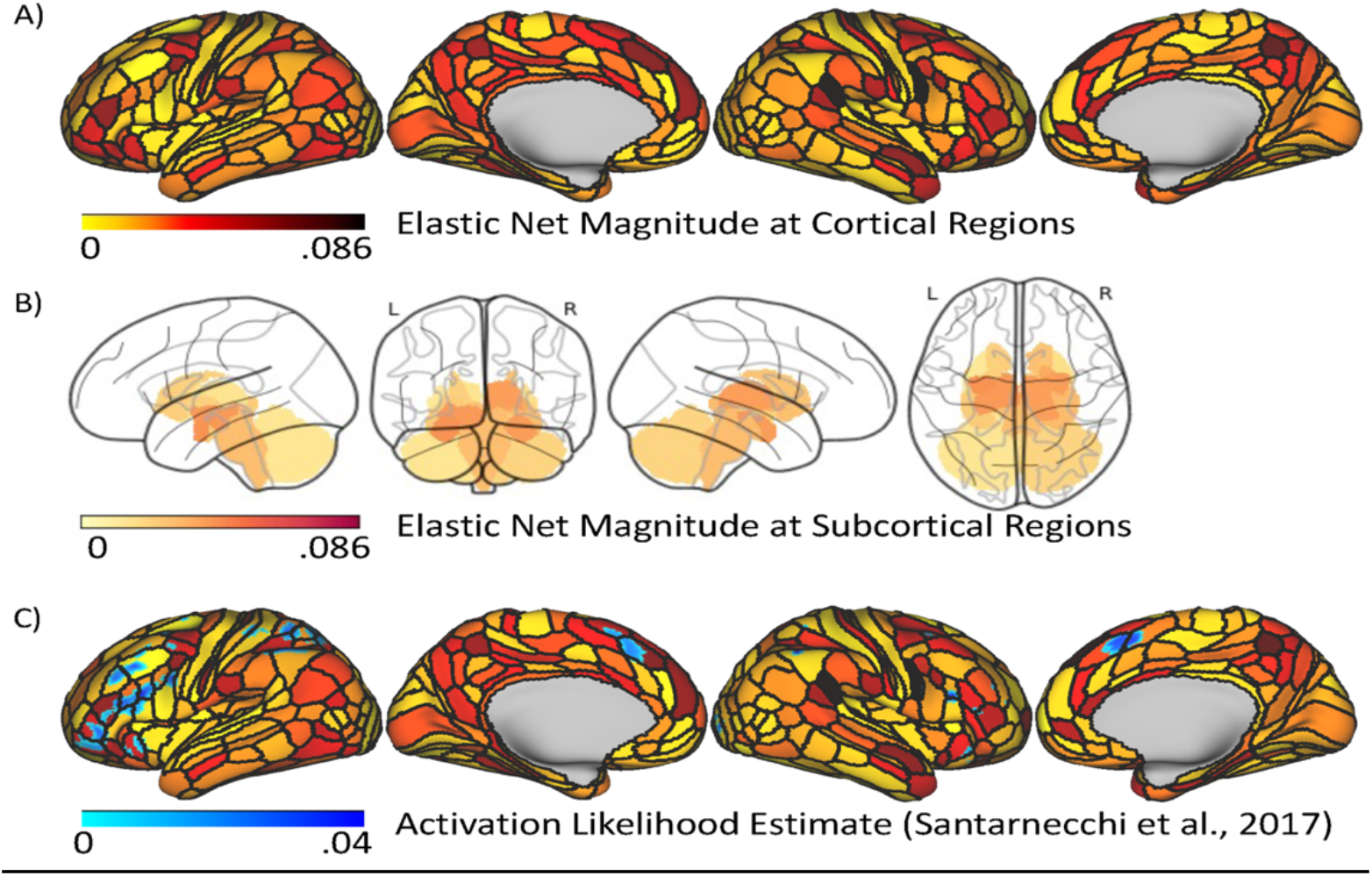
Feature importance of task-based functional MRI (tfMRI) across seven tasks. Here we combined the magnitude of Elastic Net coefficients from all seven tfMRI tasks at each brain area, weighted by the magnitude of Elastic Net coefficients of the all-task stacked model with eNet+eNet. A higher value indicates a stronger contribution to the prediction of general cognitive abilities, regardless of the directionality. 13A shows the magnitude at cortical regions while 13B shows the magnitude at subcortical regions. 13C overlays the Activation Likelihood Estimate (ALE) map of the mass-univariate associations with cognitive abilities from a previous meta-analysis (Santarnecchi et al., 2017) on top of the magnitude at the cortical regions.

**Table 1.**
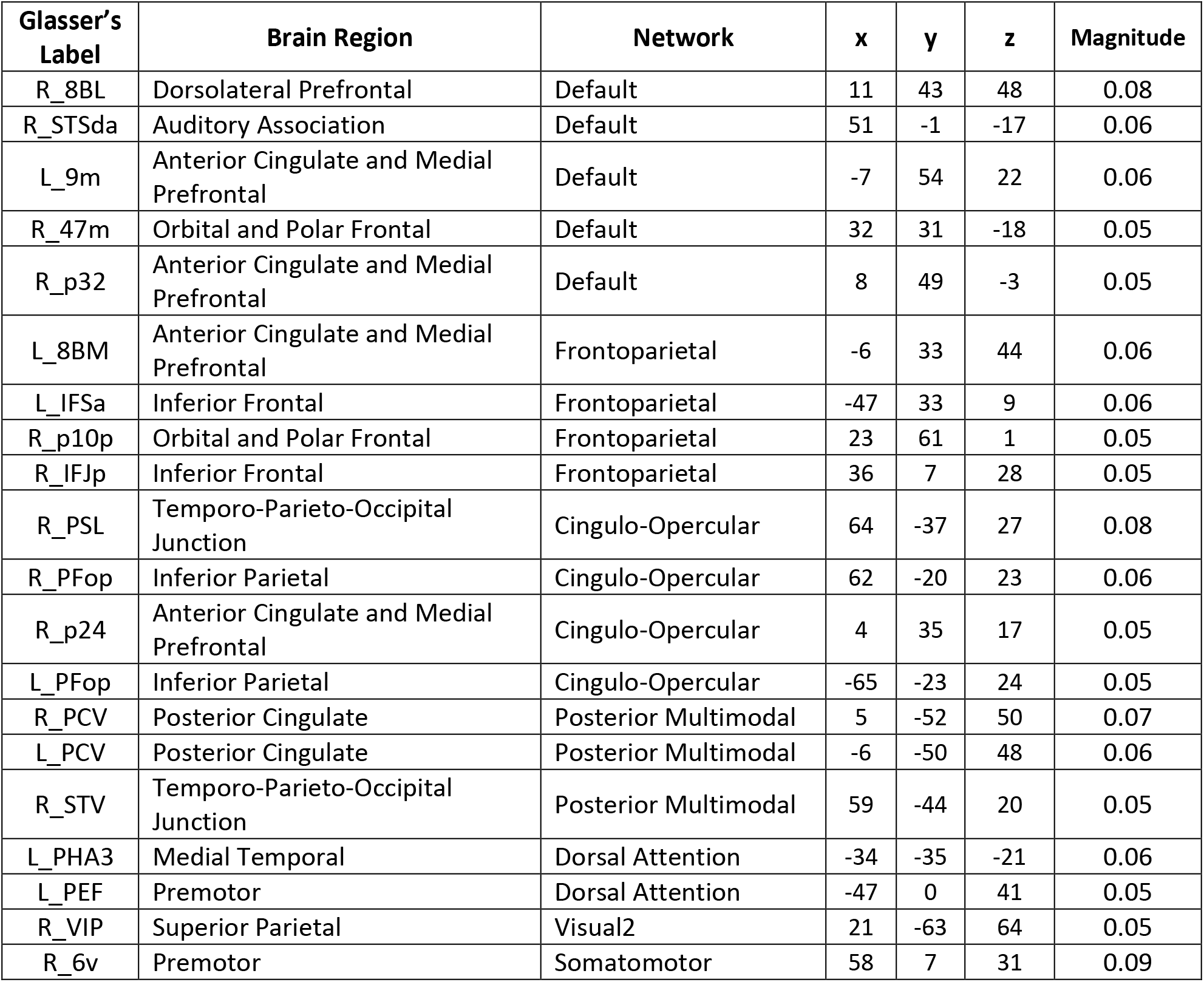
Top-20 contributing brain areas across all tfMR tasks. Here we combined the magnitude of Elastic Net coefficients from all seven tfMRI tasks, weighted by the magnitude of Elastic Net coefficients of the all-task stacked model with eNet+eNet.

| **Glasser's Label** | **Brain Region** | **Network** | **x** | **y** | **z** | **Magnitude** |
| --- | --- | --- | --- | --- | --- | --- |
| R_8BL | Dorsolateral Prefrontal | Default | 11 | 43 | 48 | 0.08 |
| R_STSda | Auditory Association | Default | 51 | -1 | -17 | 0.06 |
| L_9m | Anterior Cingulate and Medial Prefrontal | Default | -7 | 54 | 22 | 0.06 |
| R_47m | Orbital and Polar Frontal | Default | 32 | 31 | -18 | 0.05 |
| R_p32 | Anterior Cingulate and Medial Prefrontal | Default | 8 | 49 | -3 | 0.05 |
| L_8BM | Anterior Cingulate and Medial Prefrontal | Frontoparietal | -6 | 33 | 44 | 0.06 |
| L_IFSa | Inferior Frontal | Frontoparietal | -47 | 33 | 9 | 0.06 |
| R_p10p | Orbital and Polar Frontal | Frontoparietal | 23 | 61 | 1 | 0.05 |
| R_IFJp | Inferior Frontal | Frontoparietal | 36 | 7 | 28 | 0.05 |
| R_PSL | Temporo-Parieto-Occipital Junction | Cingulo-Opercular | 64 | -37 | 27 | 0.08 |
| R_PFop | Inferior Parietal | Cingulo-Opercular | 62 | -20 | 23 | 0.06 |
| R_p24 | Anterior Cingulate and Medial Prefrontal | Cingulo-Opercular | 4 | 35 | 17 | 0.05 |
| L_PFop | Inferior Parietal | Cingulo-Opercular | -65 | -23 | 24 | 0.05 |
| R_PCV | Posterior Cingulate | Posterior Multimodal | 5 | -52 | 50 | 0.07 |
| L_PCV | Posterior Cingulate | Posterior Multimodal | -6 | -50 | 48 | 0.06 |
| R_STV | Temporo-Parieto-Occipital Junction | Posterior Multimodal | 59 | -44 | 20 | 0.05 |
| L_PHA3 | Medial Temporal | Dorsal Attention | -34 | -35 | -21 | 0.06 |
| L_PEF | Premotor | Dorsal Attention | -47 | 0 | 41 | 0.05 |
| R_VIP | Superior Parietal | Visual2 | 21 | -63 | 64 | 0.05 |
| R_6v | Premotor | Somatomotor | 58 | 7 | 31 | 0.09 |

## 4. Discussion

Integrating tfMRI across tasks and with other MRI modalities via predictive modelling, we aim to boost prediction and reliability for brain MRI to capture general cognitive abilities. We found that combining tfMRI across tasks along with other MRI modalities into the all-modality stacked model gave us the best level of prediction, relative to other models, while providing an excellent test-retest reliability. Importantly, this prediction of the all-modality stacked model was primarily driven by the main three tfMRI tasks: working-memory, relational processing, and language. Combining tfMRI across tasks, especially from the three top tasks, gave us the predictive performance that was closer to the all-modality stacked model and still provided excellent test-retest reliability. This shows the importance of tfMRI from certain tasks as an information source for general cognitive abilities. Conversely, the non-task stacked model that combined sMRI and resting-state FC provided much poorer prediction. We also examined different approaches for tfMRI to be combined across tasks and with other modalities. We found 1) stacked and flat models to be comparable, and 2) a linear, non-interactive algorithm, Elastic Net, to perform well for both modality-specific and stacked models. Additionally, our use of an interpretable machinelearning algorithm, via Elastic Net, enabled us to demonstrate the crucial role of frontoparietal regions across different tfMRI tasks in predicting general cognitive abilities, in line with the parieto-frontal integration theory of intelligence (Jung & Haier, 2007).

### 4.1 Prediction of the Stacked Models

The all-modality stacked model had the highest prediction, compared to other modality-specific and stacked models, across the four measures (having the highest *r* and R^2^ and lowest MSE and MAE) and across machine-learning algorithms. This level of prediction is higher relative to those shown in other studies to date (Dubois et al., 2018; McDaniel, 2005; Mihalik et al., 2019; Pietschnig et al., 2015; Rasero et al., 2021; Sripada et al., 2020). Indeed, the level of prediction from the eNet+eNet algorithm (average across 8 CVs at *r*=.57, R^2^=.32) is much higher than the performance based on polygenic risk scores from genome-wide association (R^2^=.10) (Allegrini et al., 2019). This suggests the potential use of multimodal MRI as a robust biomarker for general cognitive abilities. Accordingly, future researchers who need a relatively high predictive and reliable brain-based biomarker for general cognitive abilities could employ our method that takes advantage of all MRI modalities available.

We found high predictive performance from the all-task and top-task stacked models that combined tfMRI from seven different and the top three tasks, respectively. This confirms the superior predictive performance of tfMRI shown in a recent study that separately investigated each tfMRI task (Sripada et al., 2020). Moreover, our results extended this task-specific work (Sripada et al., 2020), such that combining tfMRI across tasks further boosted the prediction. We also showed that, when every modality was combined into the all-modality stacked model, tfMRI from several tasks drove the prediction. This confirms that tfMRI from certain tasks provided unique and important sources of information relevant to general cognitive abilities. Next, not all tfMRI tasks drove the prediction. The top-task stacked models that included working memory, language and relational tasks already have predictive performance close to the all-task and the all-modality models. The strong performance of the working memory task, in particular, is consistent with recent studies (Pat et al., n.d.; Sripada et al., 2020). The task stacked model was also superior to the non-task stacked model even though non-task modalities (resting-state FC and sMRI) are much more commonly implemented in the literature on individual differences and cognition (Sui et al., 2020). Altogether, despite its superior performance, tfMRI has been ignored and downplayed in its importance for individual differences over non-task modalities, partly causing the unpopularity of using tfMRI as a predictive tool for cognition.

Given the popularity of resting-state FC, it might be surprising to see its poor performance for predicting general cognitive abilities here. The best performing algorithm, Elastic Net, only provided R^2^=.023 and Pearson’s *r*=.149. And this level of predictive performance is in line with other recent studies that applied nested cross-validation to predict general cognitive abilities using the Human Connectome Project (HCP) data, e.g., R^2^=.016 (Rasero et al., 2021) and R^2^=.078 and Pearson’s *r*=.26 (Sripada et al., 2020). Note though that it is possible that the HCP sample size is too small for resting-state FC to be modelled effectively. Sripada and colleagues (2021), for instance, applied a similar modelling technique they used on the HCP (Sripada et al., 2020) to a much larger study, the Adolescent Brain Cognitive Development (ABCD) Study (Casey et al., 2018) in children aged 9-10 years old (n=5,937). With the ABCD, they found much better predictive performance from resting-state FC at Pearson’s *r*=.42. Still given the differences in age ranges between the HCP and the ABCD, it is unclear if the better performance found is due to age. For instance, the predictive models for general cognitive abilities in the ABCD might capture individual differences in the brain/cognitive development that should not play a large role among young adults in the HCP. Future studies with a larger sample of young adults are needed to test this proposition.

Feature importance of the all-modality stacked model (Figure 9) also supports the important roles of certain tfMRI tasks (e.g., working-memory, relational processing, language, followed by emotion). Given Elastic Net coefficients reflect unique contributions from each feature (Zou & Hastie, 2005), these tfMRI tasks appeared to provide non-overlapping variance among themselves and with other MRI modalities in predicting the general cognitive abilities. This again reiterates the importance of including tfMRI of different tasks in the predictive model of cognitive abilities. Within the tfMRI stacked model (Figure 9), certain tasks contributed highly to the model while other tasks did not provide strong contributions (e.g., the gambling task and, to a lesser extent, the social and motor tasks). The three highly contributing tasks (working-memory, relational processing, and language) were tasks that are related to general cognitive abilities as measured by the cognitive tests (Salthouse, 2004) while other tasks (gambling, social and motor) were not. Accordingly, both the performance of the top-task stacked model and the feature importance of the all-task stacked model seems to suggest domain specificity from each task (i.e., not just any tasks, but tasks related to the target of the model).

### 4.2 Test-Retest Reliability of the Stacked Models

One of the main criticisms of tfMRI is its low test-retest reliability, compared to non-task modalities, such as sMRI (Elliott et al., 2020). Elliot and colleagues (2020) employed the same dataset (HCP) as ours and analysed test-retest reliability of tfMRI via ICC using a traditional univariate approach, i.e., separately at each prespecified region and task (Noble et al., 2021). They found poor ICC (<.4) of tfMRI signals across regions and tasks. Conversely, our predictive models drew tfMRI information across regions from the whole brain for each task. And, for the task stacked model, we further drew tfMRI information across different tasks. With this, we found ICC for certain tfMRI tasks, including language and working memory, in the good level (between .6 and .74), and more importantly, ICC for the all-task (^~^.83) and top-task (^~^.78) stacked models with enet+enet in the excellent level (Cicchetti & Sparrow, 1981). It is also noteworthy that drawing information across MRI indices using predictive modelling may improve the reliability of MRI across different modalities, not just tfMRI. Taxali and colleagues (2021), for instance, applied predictive modelling to predict cognitive abilities from resting-state FC and found ICC in the good level at around .65. These findings indicate marked improvement of test-retest reliability from predictive modelling over the classical univariate approach. Elliot and colleagues (2020) mentioned, “Without substantially higher reliability, task-fMRI measures will fail to provide biomarkers that are meaningful on an individual level.” (p. 803). Here we may have found a solution. Combining tfMRI across regions, across tasks and with MRI of other modalities gave us the best of both worlds: relatively high prediction and excellent reliability (e.g., ICC=^~^.85 for the All-Modality, enet+enet stacked model).

### 4.3 Approaches for tfMRI to be combined across tasks and with other modalities

Our study also investigated different approaches for tfMRI to be combined across tasks and with other modalities. First, we compared the stacked vs. flat models. We found similar predictive performance between the two across different machine-learning algorithms and combinations of modalities. This similarity in performance was found even though the stacked models used far fewer features in the second layers where each modality contributed to only one ‘surrogate’ feature, leaving the number of features equal to the number of modalities included in the stacked models. Accordingly, future researchers may wish to use whichever strategy, either building flat or stacked models, that may be most suitable for them. For us, we prefer the stacked models due to their strengths. First, the stacked models allowed us to demonstrate the relative contribution of each modality in predicting cognitive abilities. For instance, here we demonstrated the feature importance of the all-modality model, showing the relatively higher contribution from certain tasks. With this information, we can then focus on the modalities that account for the most variance in the target variable. This information is harder and more complicated to obtain from the flat models with much more features. Second, the stacked models enabled us to test different algorithms on their abilities to combine information from different modalities. With the flat models, an algorithm that combines features within each modality has to be the same as an algorithm that combines features across modalities. For instance, here we tested four algorithms for the second layer of our stacked models. Third, we could also take advantage of some properties of the second layer algorithms. For instance, Engemann and colleagues (2020) used Ridge regression in their first training layer, but Random Forest in their second training layer. As seen in our study, Random Forest sometimes does not result in prediction as high as other algorithms, but Engemann and colleagues (2020) and Pat and colleagues (n.d.) used Random Forest to deal with missing values from different modalities in the second training layer, allowing them to avoid dropping participants whose data from some modalities were missing.

Next, we examined different machine learning algorithms in their ability to draw information across brain features and across tfMRI tasks and MRI modalities. Over 16 combinations of algorithms, using Elastic Net both for building modality-specific models and for stacking surrogate measures across modalities (known as eNet+eNet) led to numerically highest prediction for the all-modality stacked model. In fact, despite being the only linear and additive algorithm, eNet+eNet performed significantly better than 13 out of the 16 algorithms that were non-linear and interactive. This performance of Elastic Net is in line with previous findings in resting-state FC in which a penalised regression, such as Elastic Net, usually performs on par with, if not better than, non-linear and interactive algorithms (Dadi et al., 2019). The added benefit of Elastic Net beyond predictive performance is its interpretability. Using Elastic Net coefficients as feature importance allowed us to examine which brain features, tasks and modalities contributed to the prediction of general cognitive abilities.

### 4.4 Feature Importance of tfMRI for predicting general cognitive abilities

Based on the feature importance of tfMRI, brain activity of each of the three highly contributing tasks appeared to involve similar networks, dominated by the frontoparietal and default-mode networks, and to a lesser extent, accounted for by the dorsal attention and cingulo-opercular, networks. Combining the contribution of each region across all tasks led to a distributed network of frontal and parietal brain regions that drove the prediction of general cognitive abilities. These areas include frontal regions, such as the anterior cingulate and medial prefrontal lobe, the inferior frontal lobe and the orbital and polar frontal lobe, the dorsolateral prefrontal lobe as well as parietal regions, such as the temporo-parieto-occipital junction and the inferior parietal lobe. In fact, our findings showed overlapping areas with those found in a meta-analysis of association studies (Santarnecchi et al., 2017) mainly at the frontoparietal network. This fits nicely with the parieto-frontal integration theory of the intelligence (Jung & Haier, 2007), suggesting the important role of these frontal and parietal brain regions across cognitive contexts (i.e., tfMRI tasks).

Despite the overlapping areas found between our stacking feature-importance approach and the meta-analysis (Santarnecchi et al., 2017), there are some differences in many areas, e.g., the anterior cingulate and medial prefrontal areas found in ours. Accordingly, it is important to distinguish between the two approaches. First, meta-analyses usually focus on the consistency in mass-univariate associations (e.g., between each brain region and general cognitive abilities) (Müller et al., 2018), while ours focus on relative weights in the multivariate associations (e.g., between information across brain regions and general cognitive abilities). Accordingly, our approach captures the feature importance of each brain region in the presence of other regions in the model, as opposed to ignoring the presence of other regions in the mass-univariate associations. Second, meta-analyses examine consistency in location across tfMRI tasks that predict cognitive abilities (Basten et al., 2015; Jung & Haier, 2007; Santarnecchi et al., 2017), meaning that each tfMRI task weights the same regardless of whether this task predicts cognitive abilities well. By contrast, our approach weights the importance of each task using the magnitude of Elastic Net coefficients of the task stacked model. Accordingly, our stacking feature-importance approach may contribute to the biological insights of cognitive abilities, over and above what we may have learnt from meta-analyses.

### 4.5 Implications for brain-cognition biomarkers

Beyond providing a predictive, reliable and interpretable method for capturing brain-cognition relationship, our work paves the way for developing a robust biomarker for cognitive abilities. According to the National Institute of Mental Health’s Research Domain Criteria (RDoC), cognitive abilities are considered one of the six major transdiagnostic spectrums that cut across neuropsychiatric illnesses (Morris & Cuthbert, 2012). Following the RDoC, to understand neuropsychiatric illnesses, scientists need tools to examine the transdiagnostic spectrums (such as cognitive abilities) at different units of analysis (such as gene, brain to behaviours). Recent genome-wide association studies have brought out polygenic scores that quantify cognitive abilities at the genetic level (Allegrini et al., 2019). Having a cognitive brain-based biomarker as developed in this study can serve as a link between genetics (e.g., polygenic scores) and phenotypes (e.g., cognitive abilities). Examining this link can uncover the pathway between having genetic risks to developing neuropsychiatric symptoms (Gottesman & Gould, 2003). Next, neuroscientists can also apply the brain-based biomarker to examine interventions/behaviours that may alter cognitive abilities. For instance, neuroscientists can implement the brain-based biomarker to investigate whether sleep (Taveras et al., 2017), exercise (Hötting & Röder, 2013) or extracurricular activities (Kirlic et al., 2021) improve brain processing involved in cognitive abilities, thereby deriving protective factors against many neuropsychiatric disorders. Accordingly, our biomarkers for cognitive abilities can play a vital role in the RDoC framework.

### 4.6 Limitations

Our study is not without limitations. First, to demonstrate the benefits of the task over non-task modalities, we focused on the GLM contrasts of tfMRI that reflected changes in BOLD between experimental vs. control conditions for each task. While the GLM contrasts allowed us to focus on condition-specific variance of tfMRI, we may have missed condition-non-specific variance during the tfMRI scans that may also be related to general cognitive abilities. Recent studies (Elliott et al., 2019; Greene et al., 2018) have captured condition-non-specific variance using function-connectivity during tasks and found boosted prediction and reliability over those of resting-state FC. Accordingly, future studies may further blur the line between task vs non-task modalities by including condition-non-specific function-connectivity during both tasks and rest in the stacked models and examine their performance.

Second, while the HCP (Van Essen et al., 2013) has several strengths, such as providing a large number of tfMRI tasks and high-quality, long resting-state data, the study simply does not contain sufficient numbers of participants per race/ethnicity group to be analysed. This issue is especially apparent when we compare the HCP with larger, more recent studies, such as the Adolescent Brain Cognitive Development (ABCD) Study (Casey et al., 2018) and UKBiobank (Sudlow et al., 2015). In Supplementary Materials, we listed the number of participants in each race/ethnicity group. 10 out of 14 race/ethnicity groups only included 15 participants or fewer. And two out of the four larger race/ethnicity groups only included 57 and 54 participants. This made it complicated for us to statistically control for the potential influences of race/ethnicity in the HPC. Some test folds, for instance, may have very few, if any, participants from a certain race/ethnicity group. In fact, studies using the HCP to build predictive models for cognitive abilities usually do not control for race/ethnicity (e.g., Dubois et al., 2018; Finn et al., 2015; Rasero et al., 2021; Sripada et al., 2020). We reported our attempt to do so in the Supplementary and found similar, but somewhat lower, performance after correcting for race/ethnicity. Corrections for races/ethnicities essentially assume the link between races/ethnicities, cognitive abilities and the brain. Given the insufficient numbers of participants per race/ethnicity group, the results of the correction can be biased. To avoid any over-interpretation of the link between races/ethnicities, cognitive abilities and the brain based on the HCP data, as seen in similar cases elsewhere (Fraser, 1995), we urge readers to interpret the race/ethnicity-corrected results with caution.

### 4.7 Conclusions

In conclusion, over the last decade, investigations of individual differences in the brain-cognition relationship have been dominated by non-task modalities (Sui et al., 2020). Here we show clearly that tfMRI, when used appropriately by 1) drawing information across regions from the whole brain and by 2) combining tfMRI across tasks and with other MRI modalities, can provide unique and important sources of information about individual differences in cognitive abilities. This has led to an interpretable predictive model with high prediction and excellent reliability. Our research, thus, encourages the use of tfMRI in capturing individual differences in the brain-cognition relationship for general cognitive abilities and beyond. Accordingly, future large-scale consortiums that focus on neuroimaging and individual differences should not ignore tfMRI and indeed should include tfMRI from a multitude of tasks, as pioneered by the HCP (Barch et al., 2013).

## Acknowledgements

Data were provided by the Human Connectome Project, WU-Minn Consortium (Principal Investigators: David Van Essen and Kamil Ugurbil; 1U54MH091657) funded by the 16 NIH Institutes and Centers that support the NIH Blueprint for Neuroscience Research; and by the McDonnell Center for Systems Neuroscience at Washington University. The author(s) wish to acknowledge the use of New Zealand eScience Infrastructure (NeSI) high performance computing facilities, consulting support and/or training services as part of this research. New Zealand’s national facilities are provided by NeSI and funded jointly by NeSI’s collaborator institutions and through the Ministry of Business, Innovation & Employment’s Research Infrastructure programme. URL https://www.nesi.org.nz. The authors also thank Sam Harrison and Javier Rasero for their input on data analyses. A.T. and N.P. were supported by Health Research Council Funding (21/618) and by the University of Otago. N.P. and A.S. were supported by the Intramural Research Program of the National Institute of Mental Health, National Institutes of Health, Baltimore, MD, USA (award number: ZIA-MH002957).

