## Supplementary Materials for "Capturing Brain-Cognition Relationship: Integrating Task-Based fMRI Across Tasks Markedly Boosts Prediction and Test-Retest Reliability"

**Predictive performance metrics**

- First, Pearson’s *r* is defined as

$\frac{cov(y, \hat{y})}{\sigma_{y}\sigma_{\hat{y}}} , (1)$

where *cov* is the covariance, σ is the standard deviation, *y* is the observed value and ***ŷ* is the predicted value.** Pearson’s *r* ranges from -1 to 1. The high positive Pearson’s *r* reflects high predictive accuracy, regardless of scale. Negative *r* reflects that no predictive information is present in the model.

- Second, coefficient of determination (R^2^) is defined using the sum-of-squared formulation,

$1-\frac{\sum_{i} {(\hat{y_{i}}-\bar{y} )}^{2}}{\sum_{i} {(y_{i}- \bar{y} )}^{2}}$, (2)

where *y̅* is the mean of the observed value. R^2^ is often interpreted as variance explained, with the value closer to 1 reflecting high predictive accuracy. Like Pearson’s *r,* R^2^ can be negative when there is no predictive information in the model. Note that we did not use the squared Pearson’s *r* definition of R^2^*,* which is not appropriate in the context of the out-of-sample prediction (Poldrack et al., 2020), given that it wrongly converts a negative Pearson’s *r* into a positive R^2^.

- Third, the mean square error (MSE) is defined as

$\frac{1}{n}\sum_{i=1}^{n} {(y_{i}-\hat{y_{i}})}^{2}$, (3).

MSE is sensitive to scaling and is often used to compare models across different algorithms/features. Lower MSE reflects high predictive accuracy.

- Fourth, the mean absolute error (MAE) is defined as

$\frac{1}{n}\sum_{i=1}^{n} \left| y_{i}-\hat{y_{i}} \right|, (4)$.

MAE is similar to MSE, but given the use of absolute (as opposed to squared) values, MAE can be more robust to outliers.

**Predictive performance of the combination of subcortical volume and total brain volume**

In the main analysis, we separately modelled subcortical volume and total brain volume even though both of them reflect the volume of the brain. This is due to the convention in FreeSurfer (Destrieux et al., 2010; Fischl, 2012): FreeSurfer only includes calculations from subcortical areas in the subcortical volume indices, but includes calculations from both cortical and subcortical areas in the total brain volume indices. Here we tested the predictive performance of the model that combined subcortical volume and total brain volume together. More specifically, we used Elastic Net and plotted the bootstrap distribution of the predictive performance of this combined model against the models with either subcortical volume or total brain volume by itself (see Supplementary Figure 1). We found similar performance between the combined model and the total brain volume.

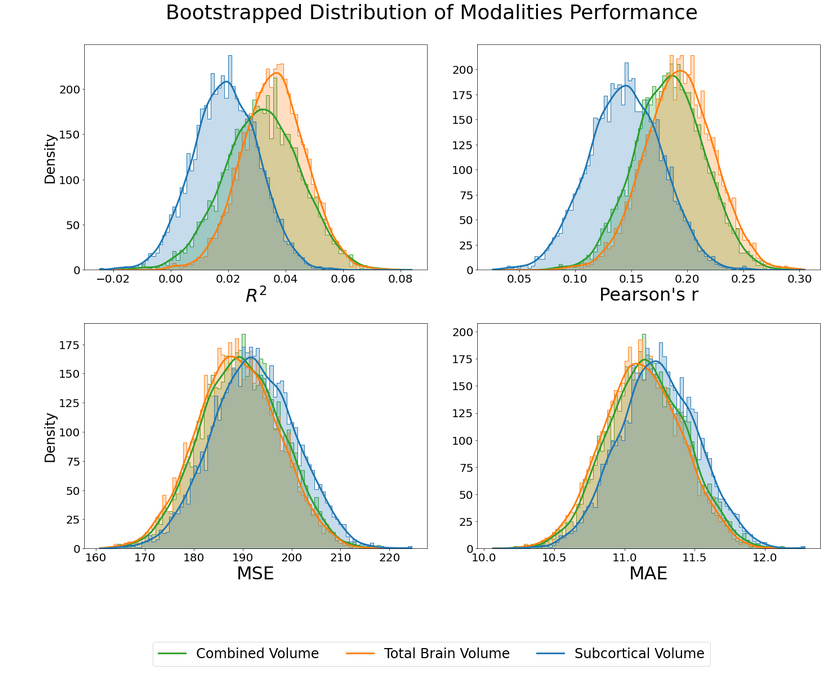

Supplementary Figure 1 The bootstrap distribution of the predictive performance of the Elastic Net model from 1) the combination of subcortical volume and total brain volume, 2) subcortical volume alone and 3) total brain volume alone. *MSE = mean square error; MAE = mean absolute error.*

**Analyses with race/ethnicity treated as another confounding variable**

In the main analysis, we treated age (Dosenbach et al., 2010; Geerligs et al., 2015), sex (Ruigrok et al., 2014; Trabzuni et al., 2013) and in-scanner movements from tfMRI and resting-state FC as confounding variables. Here, we explored the possibility to add race/ethnicity as another confounding variable.

Supplementary Table 1 shows the number of participants who identified themselves as belonging to certain race/ethnicity groups. From this table, we found that the HCP dataset is not a suitable dataset that can enable us to systematically control for potential influences of race/ethnicity. First, we found that the majority (n=606) of participants who passed our exclusion criteria (n=873) were of the same race/ethnicity, “White and not Hispanic/Latino”. Second, there were many categories of race/ethnicity that included only a few participants. For instance, only 2 people identified themselves as American Indian or Alaskan Native and not Hispanic Latino. Accordingly, controlling race/ethnicity using linear residualisation from both MRI data and cognitive abilities is likely to be problematic. Some training or test folds, for instance, may have very few, if any, participants from a certain race/ethnicity group.

Supplementary Table 1. Number of participants identified themselves as belonging to certain race/ethnicity groups.

| **Race/Ethnicity** | **n** |
| --- | --- |
| Am. Indian/Alaskan Nat._&_Not Hispanic/Latino | 2 |
| Asian/Nat. Hawaiian/Othr Pacific Is._&_Hispanic/Latino | 1 |
| Asian/Nat. Hawaiian/Othr Pacific Is._&_Not Hispanic/Latino | 54 |
| Asian/Nat. Hawaiian/Othr Pacific Is._&_Unknown or Not Reported | 2 |
| Black or African Am._&_Hispanic/Latino | 1 |
| Black or African Am._&_Not Hispanic/Latino | 117 |
| Black or African Am._&_Unknown or Not Reported | 1 |
| More than one_&_Hispanic/Latino | 7 |
| More than one_&_Not Hispanic/Latino | 15 |
| Unknown or Not Reported_&_Hispanic/Latino | 11 |
| Unknown or Not Reported_&_Not Hispanic/Latino | 2 |
| White_&_Hispanic/Latino | 57 |
| White_&_Not Hispanic/Latino | 606 |
| White_&_Unknown or Not Reported | 6 |

To mitigate the issue of having a low number of participants from certain race/ethnicity groups, we excluded the groups with fewer than 20 participants. Supplementary Table 2 below shows the four groups that were left in the analysis. With this additional exclusion criterion based on race/ethnicity, we attempted to reproduce the analysis done in the main analyses without controlling for race/ethnicity.

Supplementary Table 2. Number of participants who identified themselves as belonging to certain race/ethnicity groups after excluding the groups with fewer than 20 participants

| **Race/Ethnicity** | **n** |
| --- | --- |
| Asian/Nat. Hawaiian/Othr Pacific Is._&_Not Hispanic/Latino | 54 |
| Black or African Am._&_Not Hispanic/Latino | 117 |
| White_&_Hispanic/Latino | 57 |
| White_&_Not Hispanic/Latino | 606 |

Here we linearly residualised the dummified race/ethnicity along with age and gender from both MRI data and cognitive abilities separately from training and test data. See Supplementary Figure 2-8 for the detailed results. Briefly, with this set-up, while numerically the predictive performance for all models was lower, the all-task stacked models still performed much higher than the non-task stacked models. And we were able to reproduce most of the differences in predictive performance (e.g., regarding modalities, algorithms, stacked vs. flat models) we found without controlling for race. This suggests the robustness of the task-based fMRI data in capturing cognitive abilities.

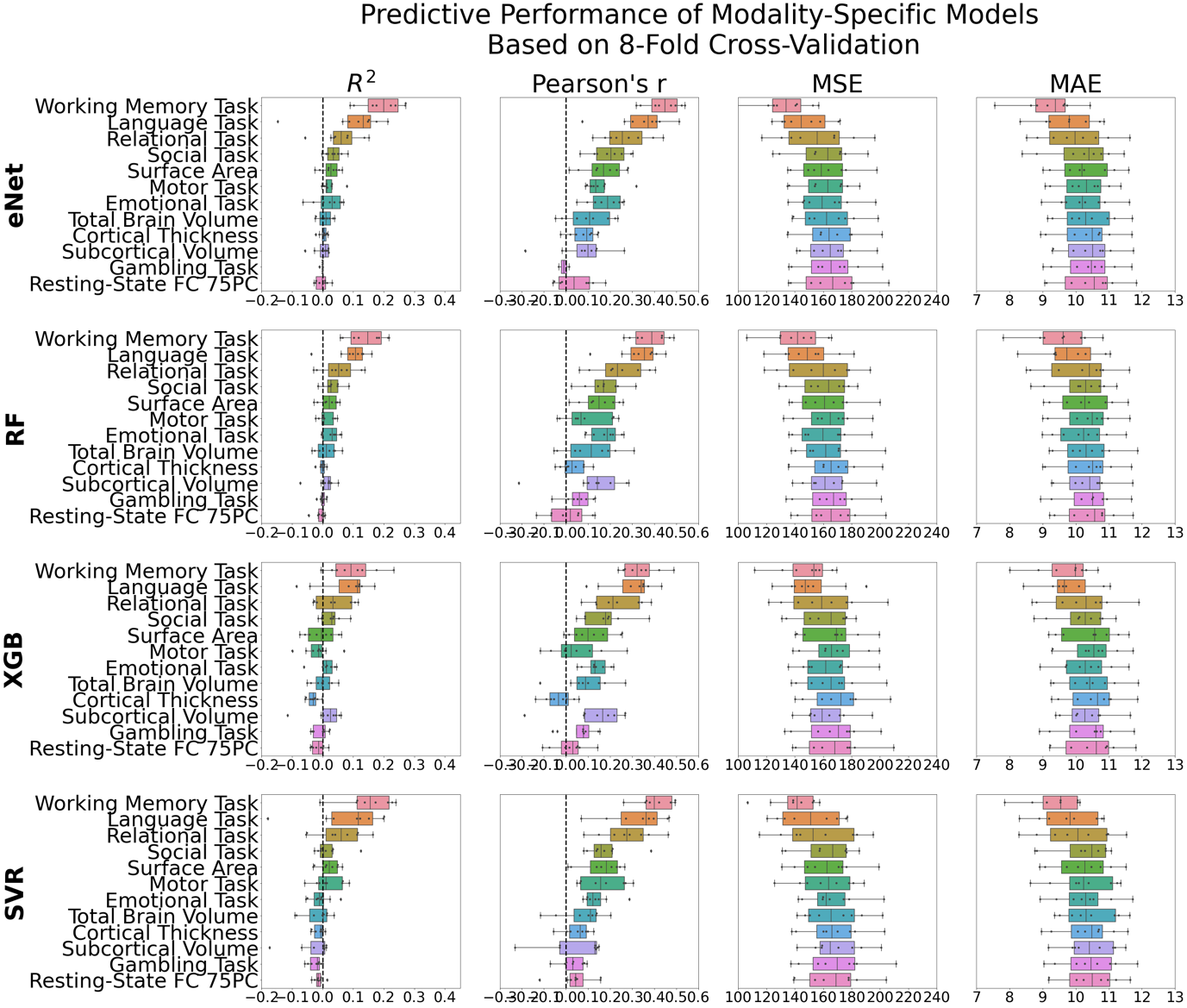

*Supplementary Figure 2. Predictive performance of modality-specific models based on the 8-fold CV across the four algorithms after controlling for race/ethnicity. Each dot represents predictive performance from each of the eight held-out folds. R^2^=coefficient of determination; eNet = Elastic Net; RF = Random Forest; XGB = XGBoost; SVR = Support Vector Regression.*

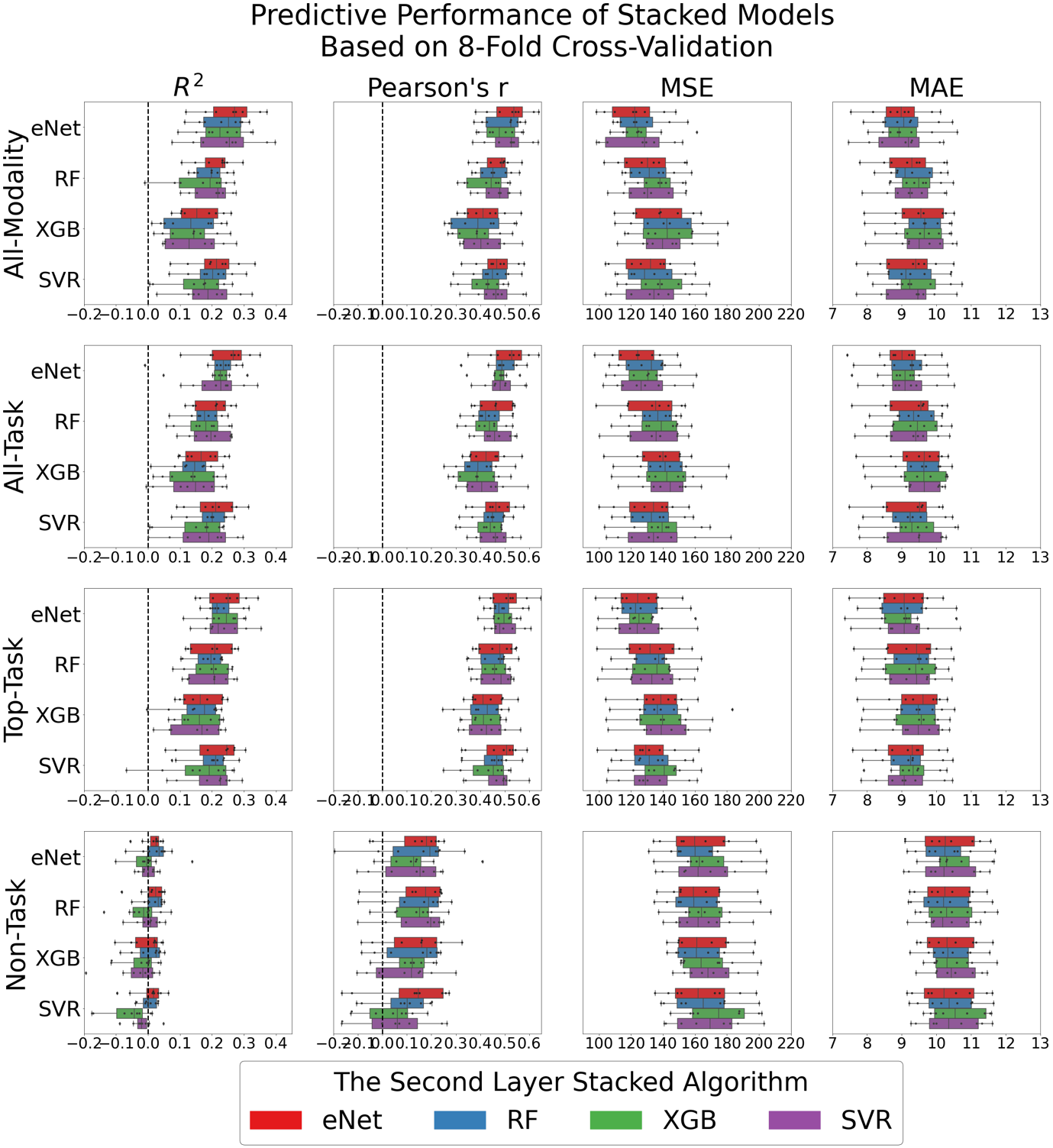

*Supplementary Figure 3. Predictive performance of stacked models based on the 8-fold CV across the four-by-four algorithms after controlling for race/ethnicity. Each dot represents predictive performance from each of the eight held-out folds. R^2^=coefficient of determination; eNet = Elastic Net; RF = Random Forest; XGB = XGBoost; SVR = Support Vector Regression.*

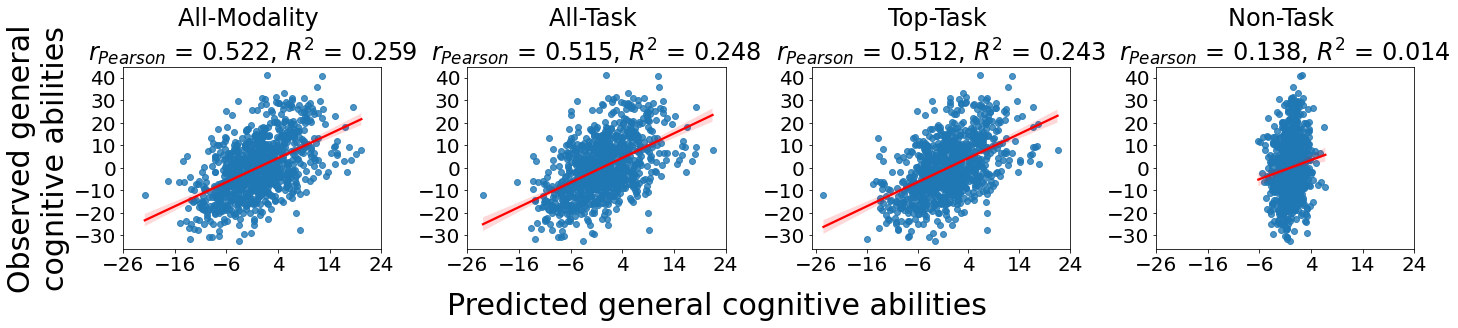

*Supplementary Figure 4. Scatter plots depicting the relationships between predicted and observed values of general cognitive abilities across the eight held-out folds of the stacked models trained on Elastic Net across both training layers after controlling for race/ethnicity. Note that Pearson’s r and R^2^ values are based on the average of the performance across 8-fold CVs.*

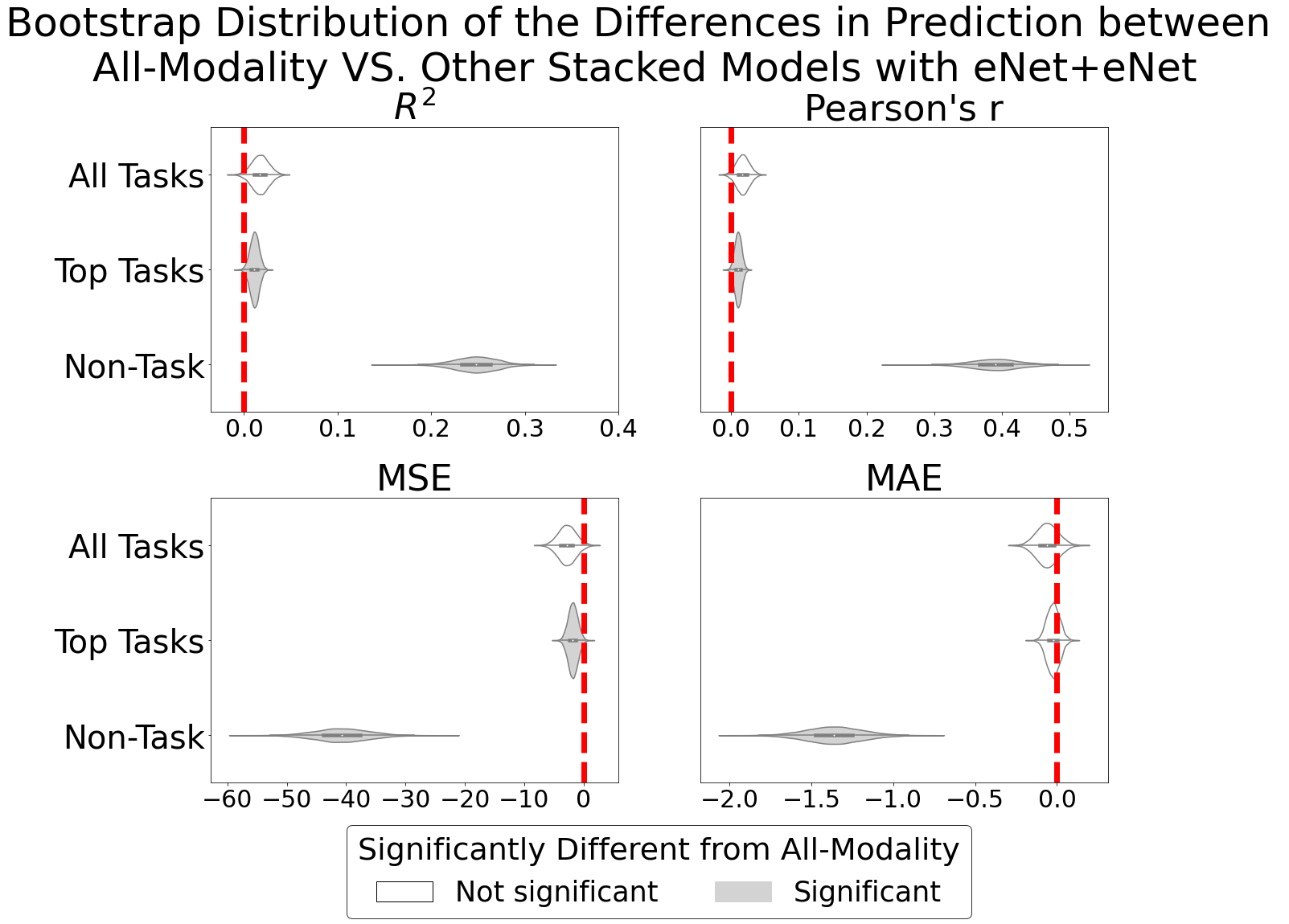

*Supplementary Figure 5. Bootstrap distribution of the differences in prediction between the all-task stacked model and other stacked models trained on Elastic Net across both training layers after controlling for race/ethnicity. For R^2^ and Pearson’s r, values lower than zero indicate better performance than the all-task stacked model. For mean square error (MSE) and mean absolute error (MAE), values lower than zero indicate worse performance than the all-task stacked model.*

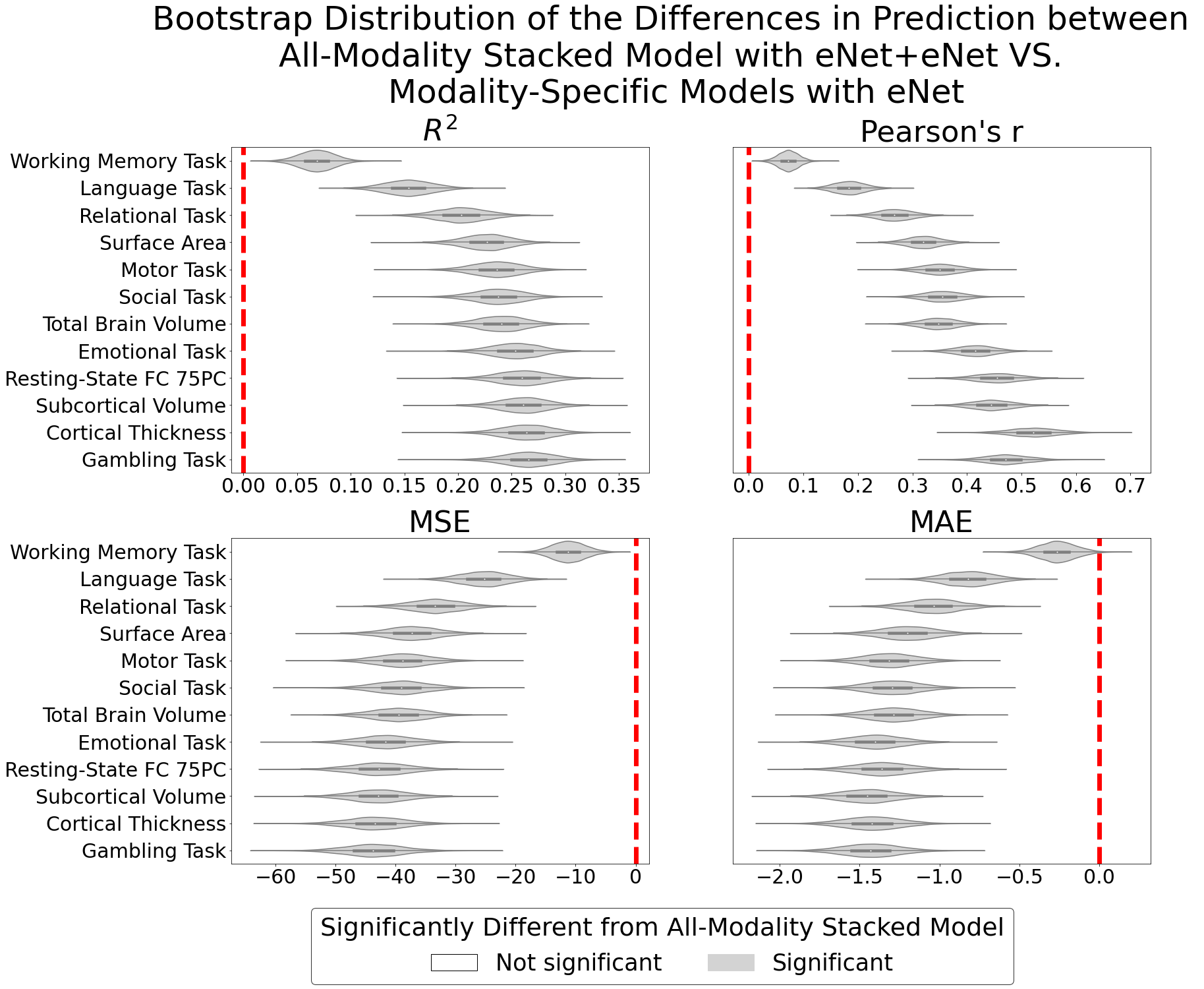

*Supplementary Figure 6. Bootstrap distribution of the differences in prediction between the all-modality stacked model trained on Elastic Net across both training layers and modality-specific models trained on Elastic Net after controlling for race/ethnicity. For R^2^ and Pearson’s r, values lower than zero indicate better performance than the all-modality stacked model. For mean square error (MSE) and mean absolute error (MAE), values lower than zero indicate worse performance than the all-modality stacked model.*

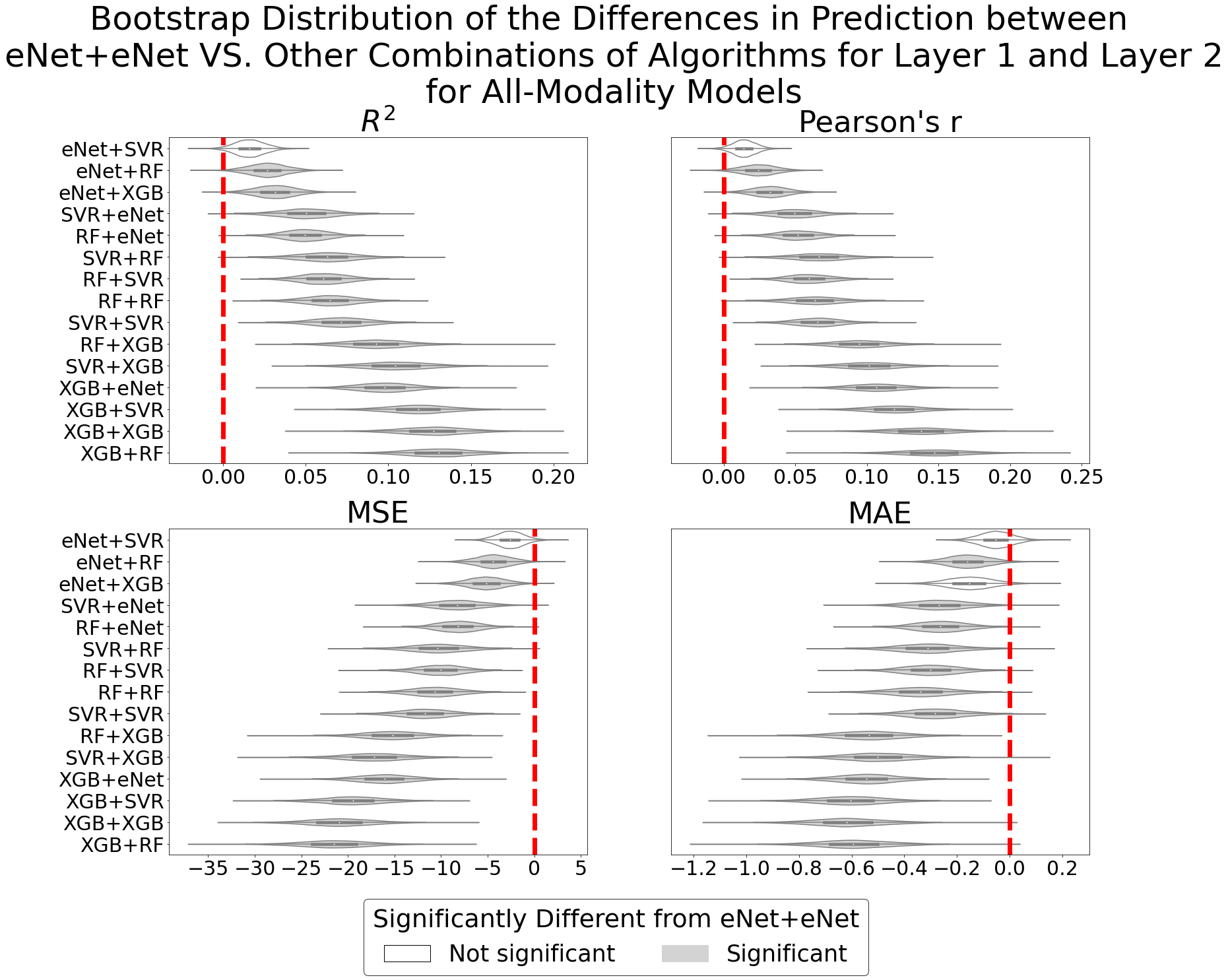

*Supplementary Figure 7. Bootstrap distribution of the differences in prediction between the all-modality stacked model trained on Elastic Net across both training layers and the all-modality stacked model trained on other combinations of algorithms after controlling for race/ethnicity. For Pearson’s r and R^2^, values lower than zero indicate better performance than the all-modality stacked model with eNet+eNet. For mean square error (MSE) and mean absolute error (MAE), values lower than zero indicate worse performance than the all-modality stacked model with eNet+eNet. The algorithm on the left of the plus symbol denotes the algorithm used for the first training layer, and the algorithm on the right of the plus symbol denotes the algorithm used for the second training layer. eNet = Elastic Net; RF = Random Forest; XGB = XGBoost; SVR = Support Vector Regression.*

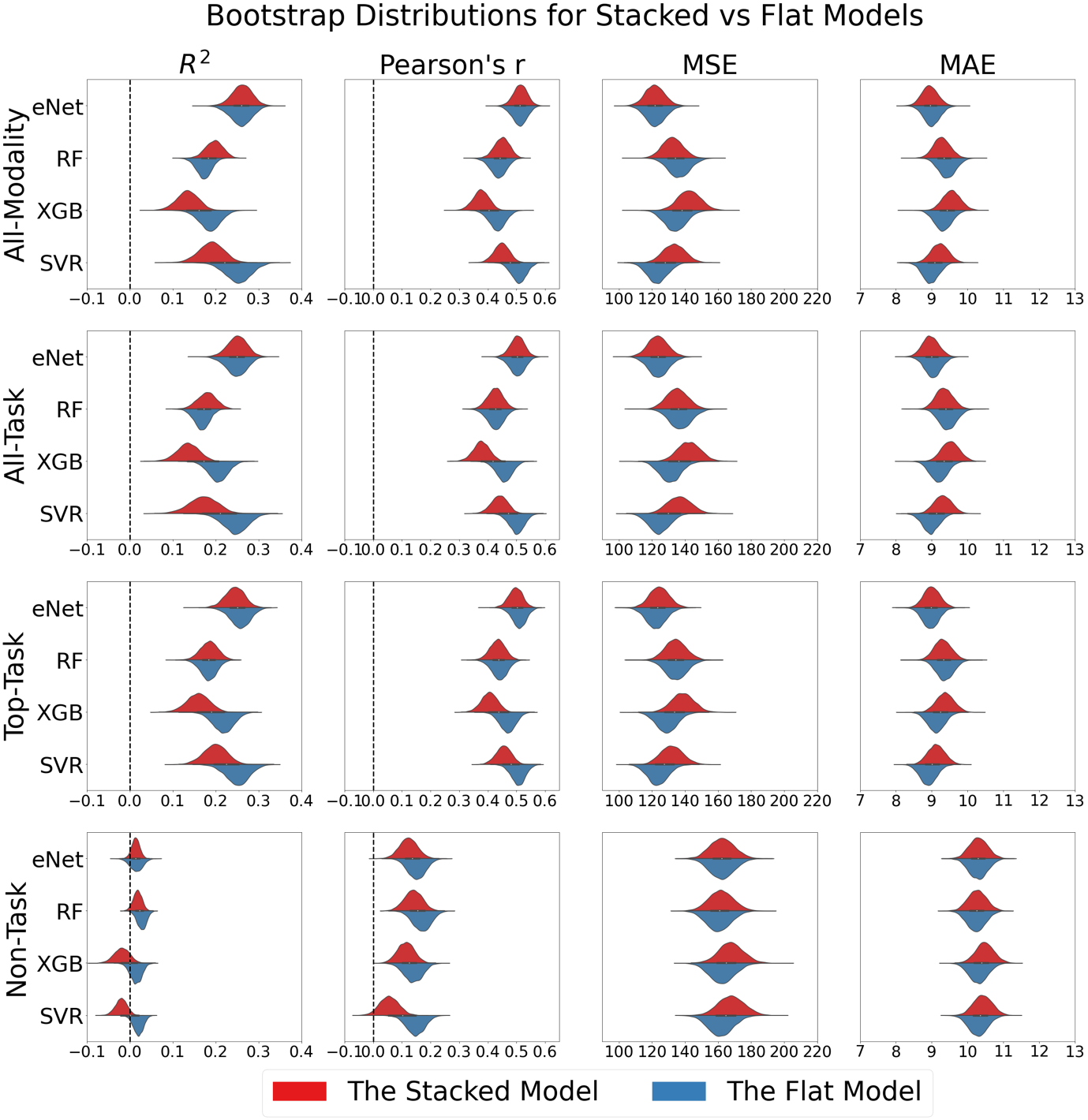

*Supplementary Figure 8. Bootstrap distribution of the predictive performance of stacked and flat models across algorithms and combinations of modalities after controlling for race/ethnicity. For the stacked models, we only plotted bootstrap distributions of models with the same machine-learning algorithms across the two training layers. eNet = Elastic Net; RF = Random Forest; XGB = XGBoost; SVR = Support Vector Regression.*

**Feature Importance for Top tfMRI Models**

*Supplementary Table 3.* *Top-20 most contributing brain regions for working memory tfMRI. The x,y,z coordinates are in MNI space.*

| **Glasser’s Label** | **Brain Region** | **Network** | **x** | **y** | **z** | **Coef**  **M** | **Coef SD** |
| --- | --- | --- | --- | --- | --- | --- | --- |
| R_33pr | Anterior Cingulate and Medial Prefrontal | Frontoparietal | 3 | 11 | 28 | 0.34 | 0.16 |
| R_7Pm | Superior Parietal | Frontoparietal | 5 | -67 | 50 | 0.32 | 0.12 |
| R_d32 | Anterior Cingulate and Medial Prefrontal | Frontoparietal | 6 | 39 | 27 | 0.29 | 0.24 |
| L_7Pm | Superior Parietal | Frontoparietal | -5 | -68 | 49 | 0.27 | 0.09 |
| R_AVI | Insular and Frontal Opercular | Frontoparietal | 32 | 25 | -4 | 0.26 | 0.09 |
| R_EC | Medial Temporal | Default | 20 | -11 | -27 | 0.31 | 0.06 |
| L_TE1a | Lateral Temporal | Default | -60 | -4 | -25 | -0.24 | 0.08 |
| L_31pv | Posterior Cingulate | Default | -10 | -44 | 33 | -0.24 | 0.09 |
| R_PHA1 | Medial Temporal | Default | 20 | -34 | -17 | -0.30 | 0.12 |
| R_8BL | Dorsolateral Prefrontal | Default | 11 | 43 | 48 | -0.47 | 0.30 |
| R_VIP | Superior Parietal | Visual2 | 21 | -63 | 64 | 0.31 | 0.08 |
| L_FST | MT+ Complex and Neighboring Visual Areas | Visual2 | -48 | -68 | 5 | 0.24 | 0.11 |
| L_V8 | Ventral Stream Visual | Visual2 | -33 | -74 | -15 | -0.26 | 0.04 |
| L_33pr | Anterior Cingulate and Medial Prefrontal | Cingulo-Opercular | -4 | 9 | 28 | 0.26 | 0.15 |
| R_IFSa | Inferior Frontal | Cingulo-Opercular | 48 | 39 | 2 | -0.33 | 0.11 |
| R_PFop | Inferior Parietal | Cingulo-Opercular | 62 | -20 | 23 | -0.41 | 0.19 |
| L_PCV | Posterior Cingulate | Posterior Multimodal | -6 | -50 | 48 | 0.38 | 0.06 |
| R_PCV | Posterior Cingulate | Posterior Multimodal | 5 | -52 | 50 | 0.28 | 0.06 |
| R_AAIC | Insular and Frontal Opercular | Orbito-Affective | 35 | 15 | -12 | 0.27 | 0.11 |
| L_AIP | Superior Parietal | Dorsal Attention | -40 | -39 | 41 | 0.32 | 0.16 |

*Supplementary Table 4.* *Top-20 most contributing brain regions for language tfMRI. The x,y,z coordinates are in MNI space.*

| **Glasser’s label** | **Brain Region** | **Network** | **x** | **y** | **z** | **Coef**  **M** | **Coef SD** |
| --- | --- | --- | --- | --- | --- | --- | --- |
| L_8BM | Anterior Cingulate and Medial Prefrontal | Frontoparietal | -6 | 33 | 44 | 0.69 | 0.44 |
| L_AVI | Insular and Frontal Opercular | Frontoparietal | -31 | 25 | -4 | 0.43 | 0.37 |
| L_a47r | Inferior Frontal | Frontoparietal | -41 | 48 | -13 | 0.30 | 0.32 |
| R_31a | Posterior Cingulate | Frontoparietal | 6 | -40 | 43 | -0.30 | 0.28 |
| R_p10p | Orbital and Polar Frontal | Frontoparietal | 23 | 61 | 1 | -0.30 | 0.22 |
| L_IFSa | Inferior Frontal | Frontoparietal | -47 | 33 | 9 | -0.58 | 0.49 |
| L_9m | Anterior Cingulate and Medial Prefrontal | Default | -7 | 54 | 22 | 0.54 | 0.53 |
| L_47l | Inferior Frontal | Default | -47 | 29 | -12 | 0.48 | 0.38 |
| R_TGd | Lateral Temporal | Default | 35 | 14 | -37 | 0.41 | 0.38 |
| L_PGi | Inferior Parietal | Default | -49 | -65 | 27 | 0.36 | 0.28 |
| L_9p | Dorsolateral Prefrontal | Default | -19 | 47 | 38 | 0.28 | 0.24 |
| L_8BL | Dorsolateral Prefrontal | Default | -11 | 38 | 52 | 0.27 | 0.19 |
| L_PFop | Inferior Parietal | Cingulo-Opercular | -65 | -23 | 24 | 0.24 | 0.08 |
| L_p24 | Anterior Cingulate and Medial Prefrontal | Cingulo-Opercular | -5 | 37 | 13 | -0.25 | 0.23 |
| R_PSL | Temporo-Parieto-Occipital Junction | Cingulo-Opercular | 64 | -37 | 27 | -0.69 | 0.47 |
| R_6v | Premotor | Somatomotor | 58 | 7 | 31 | 0.75 | 0.50 |
| R_1 | Somatosensory and Motor | Somatomotor | 48 | -22 | 54 | -0.34 | 0.49 |
|  | Thalamus right | subcortex |  |  |  | 0.28 | 0.31 |
| R_PCV | Posterior Cingulate | Posterior Multimodal | 5 | -52 | 50 | -0.34 | 0.49 |
| L_PHA3 | Medial Temporal | Dorsal Attention | -34 | -35 | -21 | -0.53 | 0.42 |

*Supplementary Table 5.* *Top-20 most contributing brain regions for relational tfMRI. The x,y,z coordinates are in MNI space.*

| **Glasser’s Label** | **Brain Region** | **Network** | **x** | **y** | **z** | **Coef**  **M** | **CoefSD** |
| --- | --- | --- | --- | --- | --- | --- | --- |
| R_47m | Orbital and Polar Frontal | Default | 32 | 31 | -18 | -0.15 | 0.10 |
| R_POS1 | Posterior Cingulate | Default | 11 | -57 | 15 | -0.16 | 0.07 |
| R_47l | Inferior Frontal | Default | 44 | 32 | -15 | -0.17 | 0.11 |
| L_47m | Orbital and Polar Frontal | Default | -37 | 31 | -17 | -0.20 | 0.13 |
| L_FST | MT+ Complex and Neighboring Visual Areas | Visual2 | -48 | -68 | 5 | 0.19 | 0.12 |
| R_VIP | Superior Parietal | Visual2 | 21 | -63 | 64 | 0.18 | 0.11 |
| L_PIT | Ventral Stream Visual | Visual2 | -47 | -77 | -11 | -0.16 | 0.08 |
| L_55b | Premotor | Language | -49 | -1 | 50 | 0.17 | 0.09 |
| L_SFL | Dorsolateral Prefrontal | Language | -8 | 17 | 63 | 0.15 | 0.06 |
| R_45 | Inferior Frontal | Language | 50 | 27 | 0 | -0.18 | 0.11 |
| R_p9-46v | Dorsolateral Prefrontal | Frontoparietal | 46 | 33 | 25 | 0.18 | 0.10 |
| R_IFJp | Inferior Frontal | Frontoparietal | 36 | 7 | 28 | 0.17 | 0.11 |
| R_i6-8 | Dorsolateral Prefrontal | Frontoparietal | 32 | 9 | 57 | 0.15 | 0.09 |
|  | Diencephalon ventral | subcortex |  |  |  | 0.18 | 0.09 |
| L_ProS | Posterior Cingulate | Visual1 | -23 | -55 | 3 | 0.15 | 0.10 |
| L_FOP2 | Insular and Frontal Opercular | Somatomotor | -44 | -5 | 14 | -0.17 | 0.11 |
| R_TPOJ2 | Temporo-Parieto-Occipital Junction | Posterior Multimodal | 56 | -57 | 10 | -0.15 | 0.08 |
| R_AAIC | Insular and Frontal Opercular | Orbito-Affective | 35 | 15 | -12 | 0.15 | 0.08 |
| R_6a | Premotor | Dorsal Attention | 26 | -2 | 53 | 0.16 | 0.08 |
| R_TA2 | Auditory Association | Auditory | 50 | 4 | -8 | -0.16 | 0.07 |
